# Characterising replisome disassembly in human cells

**DOI:** 10.1101/2022.07.12.499744

**Authors:** Rebecca M. Jones, Joaquin Herrero Ruiz, Shaun Scaramuzza, Sarmi Nath, Marta Henklewska, Toyoaki Natsume, Francisco Romero, Masato T. Kanemaki, Agnieszka Gambus

## Abstract

To ensure faultless duplication of the entire genome, eukaryotic replication initiates from thousands of replication origins. Replication forks emanating from origins move through the chromatin until they encounter forks from neighbouring origins, at which point they terminate. In the final stages of this process the replication machinery (replisome) is unloaded from chromatin and disassembled. Work from model organisms has elucidated that during replisome unloading, the MCM7 subunit of the terminated replicative helicase is polyubiquitylated and processed by p97/VCP segregase, leading to disassembly of the helicase and the replisome, which is built around it. In higher eukaryotes (worms, frogs, mouse embryonic stem cells), MCM7 ubiquitylation is driven by a Cullin2-based ubiquitin ligase, with LRR1 as a substrate receptor. To date, most of our knowledge of replication termination comes from model organisms and embryonic systems and little is known about how this process is executed and regulated in human somatic cells. Here we thus established methods to study replisome disassembly in human model cell lines. Using flow cytometry, immunofluorescence microscopy and chromatin isolation with western blotting, we can visualise unloading of the replisome (MCM7 and CDC45) from chromatin by the end of S-phase. We observe interaction of replicative helicase (CMG complex) with CUL2^LRR1^ and ubiquitylation of MCM7 on chromatin, specifically in S-phase, suggesting that this is a replication-dependent modification. Importantly, we are able to show that replisome disassembly in this system also requires Cullin2, LRR1 and p97, demonstrating conservation of the mechanism. Moreover, we present evidence that the back-up mitotic replisome disassembly pathway is also recapitulated in human somatic cells. Finally, while we find that treatment with small molecule inhibitors against cullin-based ubiquitin ligases (CULi) and p97 (p97i) does lead to phenotypes of replisome disassembly defects, they also both lead to induction of replication stress in somatic cells, which limits their usefulness as tools to specifically target replisome disassembly processes in this setting.

## Introduction

Cell division is the basis for the propagation of life and requires accurate duplication of all genetic information. The perfect execution of DNA replication is essential to maintain a stable genome and protect from diseases such as genetic disorders, cancer and premature ageing (1). Fundamental studies over the last 50 years have led to a step-change in our understanding of DNA replication initiation and DNA synthesis (2), but until the discovery of the first elements of the replisome disassembly mechanism in 2014, the termination stage of eukaryotic replication was mostly unexplored. DNA replication termination is triggered whenever two replication forks, coming from neighboring origins, meet each other and converge in a head-to-head orientation. During termination, the final fragments of DNA need to be replicated, the replisomes disassembled and DNA intertwines resolved. A major difficulty when investigating replication termination is that the genomic position of replication fork termination and convergence is stochastic, occurring wherever two forks meet. Throughout S-phase in eukaryotic cells, replication forks undergo initiation, elongation and termination at different times; early replication forks will terminate before late replicating forks are initiated. Therefore, it is difficult to study the termination stage of DNA replication at the ensemble level. Consequently, replication termination has thus far been predominantly studied in reconstituted biochemical systems and model organisms that allow the termination of forks in a synchronized and controlled fashion. In the last eight years, biochemistry and cell biology research carried out using *Saccharomyces cerevisiae*, *Xenopus laevis* egg extracts and *Caenorhabditis elegans* embryos have allowed us to elucidate the first model of the mechanism of Eukaryotic DNA replication termination, with the unloading of the replication machinery (replisome disassembly) being the best understood stage of this process thus far (Figure 1).

**Figure 1.**
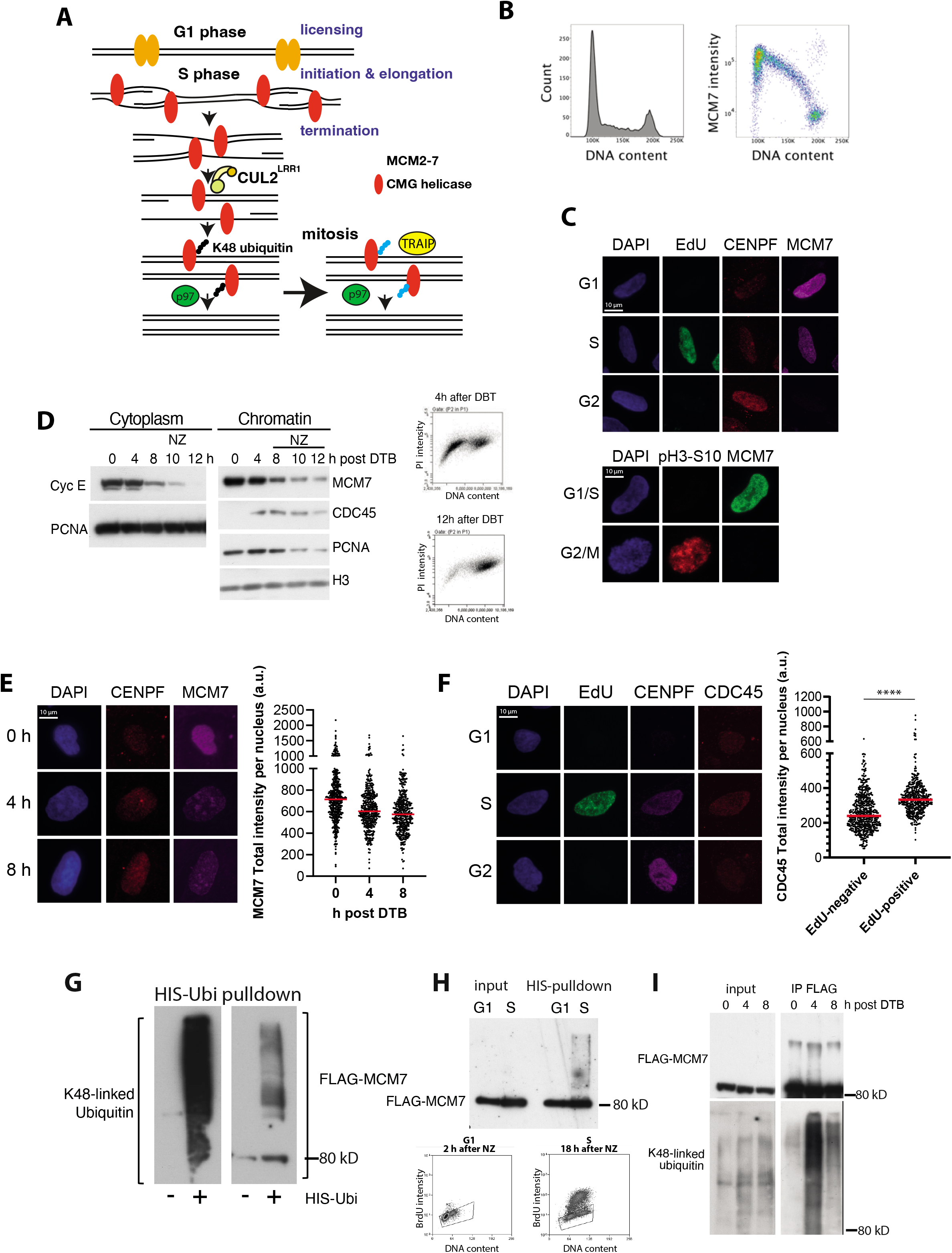
Visualisation of replisome unloading from chromatin in human somatic cells. **(A)** Schematic representing the known pathway of replisome unloading. **(B)** FACS analysis of CSK buffer extracted nuclei shows MCM7 loading and unloading from chromatin during the cell cycle. **(C)** Immunofluorescence of MCM& chromatin binding. Representative immunofluorescence images of EdU, CENPF or pH3-S10 and MCM7 binding to chromatin throughout the cell cycle. Asynchronous U2OS cells were pulse-labelled with EdU for 20 min, before cells were extracted with CSK buffer and fixed with PFA. Cells were then immunostained indicated antibodies. **(D)** Chromatin isolation and western blotting to visualise loading and unloading of CMG factors from chromatin throughout the cell cycle. U2OS cells were synchronised with double thymidine block and released for indicated length of time. Nocodazole was added at later timepoints to stop cells from entering next cell cycle. Cells were extracted with CSK buffer and chromatin samples analysed through western blotting with indicated antibodies. **(E)** Unloading of MCM7 from chromatin by immunofluorescence in arrested cells. Immunofluorescence quantification of MCM7 intensity in U2OS cells synchronised with double thymidine block and released for indicated length of time. Red lines indicate the mean. **(F)** CDC45 chromatin binding throughout the cell cycle. Representative immunofluorescence images of EdU, CENPF and CDC45 binding with chromatin throughout the cell cycle in asynchronous U2OS cells, with quantification of CDC45 intensity in EdU-negative vs EdU-positive cells (n=3). Red lines indicate the mean (p=<0.0001). **(G)** MCM7 is ubiquitylated on chromatin. U2OS cells expressing FLAG-MCM7 were optionally transfected HIS-Ubi plasmid for 48 h. Cells from asynchronous population were extracted with CSK buffer and HIS-tagged/ubiquitylated proteins were then isolated from chromatin samples and samples analysed through western blotting with indicated antibodies. **(H)** MCM7 is ubiquitylated on chromatin during S-phase. As in (G) but cells were initially synchronised with nocodazole and released. Cells were harvested at G1 and S-phase. **(I)** MCM7 is ubiquitylated on chromatin in S-phase with K48-linked ubiquitin chains. U2OS cells expressing FLAG-MCM7 were synchronised with DTB and then released for indicated time before cells were extracted with CSK buffer in denaturing conditions. FLAG-MCM7 was then isolated from chromatin with anti-FLAG beads and samples analysed through western blotting with indicated antibodies.

The eukaryotic replicative helicase is a complex formed of three components: CDC45, MCM2-7 hexamer and GINS complex (SLD5, PSF1-3), altogether referred to as the CMG complex (from the first letters of the components). Upon termination, MCM7 is specifically polyubiquitylated to promote the removal of the helicase from chromatin in a p97/VCP segregase-dependent manner. As the CMG helicase forms the organising centre of the replisome, its removal leads to disassembly of the entire replisome (3). Importantly, the main elements of this process are evolutionarily conserved in *S. cerevisiae* (4), *C. elegans* (5) and *X. laevis* egg extract (3). Following on from these initial findings, we and others went on to identify the CUL2^LRR1^ ubiquitin ligase (LRR1 is a substrate specific receptor for the ligase) as the enzyme responsible for MCM7 ubiquitylation in S-phase in higher eukaryotes (5, 6). CUL2^LRR1^ can specifically recognise terminated helicases, as its substrate binding requires availability of an interaction surface on the CMG helicase, which is obscured by the excluded lagging strand ssDNA protruding from CMG during unwinding at replication forks (7–9). Once MCM7 is ubiquitylated with K48-linked ubiquitin chains, it is recognised by the p97-UFD1-NPL4 complex with the help of the UBXN7 cofactor (3, 5, 10). Consequently, p97-driven unfolding of MCM7 leads to breaking of the CMG complex and its unloading from chromatin. Finally, when the LRR1-dependent mechanism fails, there is a back-up pathway of replisome disassembly, activated in mitosis, and regulated by the TRAIP ubiquitin ligase (5, 11). Importantly, TRAIP ubiquitin ligase was also shown to act during S-phase to unload replisomes that converge at inter-strand crosslinks (ICL) to allow for their repair (12). Much of this mechanism of replisome disassembly has been also recently confirmed to be conserved in mouse embryonic stem cells where, unlike in human cells, TRAIP is not essential for cell viability (13). Moreover, although the basic elements of this process are conserved between Fungi and Metazoa, some of the factors and regulations involved differ. For example, in *S. cerevisiae,* Mcm7 is ubiquitylated during termination by SCF^Dia2^ and TRAIP ubiquitin ligase simply does not exist. Hence, the failure of unloading during S-phase results in replisome persistence on chromatin until the next cell cycle (4).

Importantly, in higher eukaryotes, the process of replisome disassembly has been studied thus far mostly in embryonic systems; as these models pose advantages that make them particularly useful for the study of DNA replication. However, it is known that embryonic cell cycles and DNA replication are regulated somewhat differently than in somatic cells. For example, the stabilisation of CDT1 during DNA replication is not enough to induce re-replication in *Xenopus laevis* egg extract (14), as it is in human somatic cells (15); whilst origin firing is regulated by both ATM and ATR in embryonic stem cells, but mainly by ATR in somatic cells (16). Moreover, embryonic systems are known to maintain fast cell cycles with non-existent or short gap phases, with reduced regulation of cell cycle phase transition and accumulation of high levels of replication factors to sustain this fast proliferation rate. The consequences of dysregulating the process of replisome disassembly may therefore differ between the embryonic systems and somatic cells. Indeed, the first study focusing on this process in human retinoblastoma RPE1 cells has suggested that short term CRISPR mediated LRR1 depletion leads to problems with replisome disassembly during S-phase, resulting in the failure of S-phase completion due to a reduction in the pool of available CDC45 and GINS required to fire late origins (17). Finally, earlier studies of human CUL2^LRR1^ showed that it is needed to regulate the cytoplasmic pool of CDK inhibitor p21 and regulate cell migration (18).

Here, we have set out to explore the mechanism of replisome disassembly in human somatic cells. We aimed to set up assays to study this process in model cell lines, validate tools to study this process and uncover similarities and differences between somatic and embryonic regulation of replisome disassembly. We found that all elements of this process, previously reported in higher eukaryotic model systems, are conserved in human cell lines: MCM7 is ubiquitylated with K48-linked ubiquitin chains on S-phase chromatin; CUL2^LRR1^ interacts with replisome in S-phase and is required for replisome disassembly during S-phase; there is a mitotic back-up pathway of replisome disassembly and p97 is needed for unloading of replisomes both in S-phase and in mitosis. We have found however that replication progression and replisome disassembly is much more tightly regulated in tissue culture somatic cells, compared with model systems used previously to study replisome disassembly. As a result, we find that the inhibitors commonly used as tools to study this process in model systems are less appropriate for use in somatic cells.

## Results

### Replisome unloading in human cell lines

To determine the mechanism of replisome unloading from chromatin in human cells, we first established methods to visualise unloading of the replicative helicase. The main component of the replicative helicase, the MCM2-7 complex, is initially loaded onto origins of replication in late mitosis and throughout G1 stage of the cell cycle. In S-phase some of the MCM2-7 hexamers are activated to form CMGs and replisomes. During the process of DNA replication, the replisomes, as well as inactive MCM2-7 complexes (dormant origins), are unloaded as termination events occur (19, 20). Finally, by G2/M phase, all replisomes are disassembled prior to chromosome segregation. We can observe this pattern of chromatin binding of the MCM2-7 complex using FACS sorting on detergent extracted HCT116 cells, where only chromatin bound MCM7 signal is analysed. We have seen that the highest levels of chromatin-bound MCM7 are reached in G1 cells, and that these levels decrease throughout S-phase, with the majority of G2/M stage cells then having the lowest MCM7 intensities (Figure 1B). We can also visualise MCM2-7 complexes on chromatin by immunofluorescence of extracted cell nuclei and with this we can detect cells with varying levels of MCM7 on chromatin in an asynchronous population. To assign cell cycle stage status to each cell we used EdU incorporation into DNA to identify actively replicating cells; to identify G2 cells we utilised staining with CENPF/Mitosin - a large kinetochore-associated protein that is commonly used as a cell cycle marker due to its homogeneous nuclear distribution in G2 and mitosis (21); and to identify mitotic cells staining with phospho-histone H3 S10 (pH3-S10) - a universal marker of chromosome condensation in eukaryotes (22). Cells, which are negative for EdU and CENPF signal were classed as in G1. Cells positive for EdU and had increasing staining for CENPF were classed as early/mid/late S-phase. Those negative for EdU with strong staining for CENPF were classed as G2. Finally, those with strong staining for phospho-histone H3 S10 (pH3-S10) and displaying chromatin condensation phenotype by DAPI staining were classed as mitotic. In both S-phase and G2 cells, the level of MCM7 on chromatin was lower than that seen in G1 cells (Figure 1C). To further support this, U2OS cells were synchronised in early S-phase with a double thymidine block (DTB) and then released into S-phase, following which we analysed the levels of chromatin-bound MCM7 through western blotting (Figure 1D) and immunofluorescence (Figure 1E). In both assays, we could detect the highest levels of MCM7 at the 0 h time point (early S-phase), with levels dropping throughout S-phase. In order to detect the active CMG complex specifically, we also analysed levels of chromatin-bound CDC45 through western blotting (Figure 1D) and immunofluorescence (Figure 1F). As expected, we could only detect CDC45 associating with chromatin during S-phase when DNA replication is taking place. With all this, we had now established conditions for analysing replisome disassembly in human cells.

### MCM7 ubiquitylation

Previous studies into the mechanism of replication termination revealed that MCM7 is ubiquitylated prior to replisome disassembly (3, 4). We next aimed to observe the ubiquitylation of MCM7 on chromatin. To this end, we established a U2OS cell line expressing FLAG-tagged MCM7 (FLAG-MCM7), as well as one expressing FLAG-MCM4 as a control. Expression levels of this exogenous FLAG-MCM7 are very close to those of the endogenous MCM7, and FLAG-MCM7 binds chromatin equally well (Supp Figure 1A). Cells expressing FLAG-MCM7 also show a normal cell cycle profile (Supp Figure 1B) and display a similar profile of FLAG-MCM7 unloading from chromatin, compared to endogenous MCM7 (Supp Figure 1C). Moreover, FLAG-MCM7 co-immunoprecipitates with the CMG helicase component GINS (Supp Figure 1D). Importantly, the immunoblotting antibody signal is much cleaner for FLAG than endogenous MCM7 antibody, which allows us to better detect ubiquitylated forms of FLAG-MCM7. To analyse this ubiquitylated MCM7, we transfected cells with a HIS-tagged ubiquitin plasmid and performed a HIS pull-down to isolate all ubiquitylated proteins. MCM7 has been reported previously to be ubiquitylated on chromatin in human cells (23) and we could reproduce this observation (Figure 1G). To confirm that this ubiquitylation occurs in S-phase, cells were synchronised in mitosis with nocodazole and released into G1 and then S-phase. Using the HIS pull-down assay, we could clearly see that FLAG-MCM7 is indeed ubiquitylated specifically in S-phase and not in the G1 stage of the cell cycle (Figure 1H), despite higher levels of MCM7 on G1 chromatin (Figure 1B). To further support this, we immunoprecipitated FLAG-MCM7 from chromatin of cells synchronised in S-phase with release from double thymidine block and could detect a ladder of bands over FLAG-MCM7 at the 4 h time-point, post release (Figure 1I), suggesting that it is ubiquitylated upon recovery of DNA synthesis (Supp Figure 1E). Moreover, using an antibody specific to K48-linked ubiquitin chains, we could detect a strong signal over the size of FLAG-MCM7, supporting the model that MCM7 is ubiquitylated in S-phase with K48-linked ubiquitin chains (Figure 1I).

### CUL2^LRR1^ interacts with the replisome on chromatin in S-phase

Having established methods to observe replisome disassembly and ubiquitylation of MCM7 during this process, we next set out to analyse the mechanism of this process. First, we wanted to determine whether the CUL2^LRR1^ ubiquitin ligase interacts with the replisome during S-phase. Such interactions would support the model that CUL2^LRR1^ drives the ubiquitylation and unloading of the replisome at termination, suggested by phenotypes of CRISPR-Cas9 guided knockout of LRR1 in RPE1 cells (17). To do this, we transiently expressed FLAG-tagged LRR1 in U2OS cells and confirmed that it can interact with endogenous CUL2 through a FLAG IP (Supp Fig 2A). Cells overexpressing this FLAG-LRR1 did not exhibit cell viability defects (Supp Fig 2B), nor any differences to cell cycle and S-phase progression (Supp Fig 2C and D). Following this, we confirmed that we could observe co-immunoprecipitation of the replisome components: MCM7, CDC45 and GINS, with FLAG-LRR1 from S-phase chromatin (Figure 2A).

**Figure 2.**
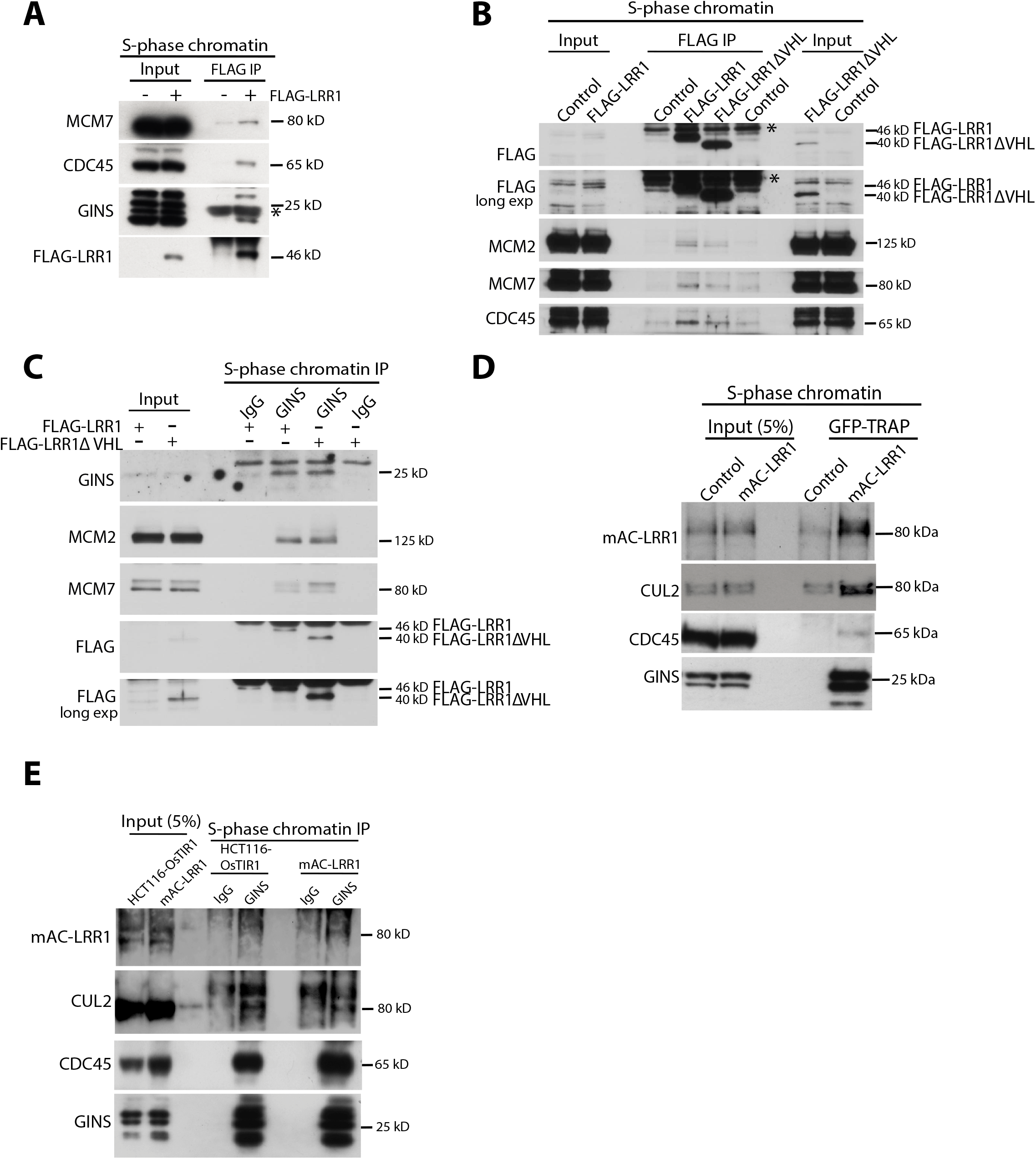
CUL2^LRR1^ interacts with the replisome during S-phase. **(A)** LRR1 interacts with replisome components on chromatin during S-phase. Chromatin extracts of FLAG-LRR1 expressing cells synchronised in S-phase by DTB were immunoprecipitated with FLAG M2 beads. Immunoprecipitated samples were analysed by western blotting with antibodies against the indicated proteins. **(B)** LRR1 mutant unable to interact with CUL2 can still interact with the replisome. Chromatin extracts of FLAG-LRR1 or FLAG-LRR1ΔVHL expressing cells synchronised in S-phase (DTB) were immunoprecipitated with FLAG-M2 beads. Immunoprecipitated samples were analysed by western blotting with indicated antibodies. **(C)** LRR1 mutant unable to interact with CUL2 can still interact with the replisome. GINS was immunoprecipitated from chromatin extracts of FLAG-LRR1 or FLAG-LRR1ΔVHL expressing cells synchronised in S-phase (DTB). Immunoprecipitated samples were analysed by western blotting with indicated antibodies. **(D)** Endogenous mAC-tagged LRR1 can interact with replisome components on chromatin in S-phase. Chromatin extracts of mAC-LRR1 expressing cells synchronised in S-phase (with lovastatin release) were enriched on GFP-Trap magnetic agarose. Immunoprecipitated samples were analysed by western blotting with indicated antibodies. **(E)** CUL2 and endogenous LRR1 can interact with GINS on S-phase chromatin. Chromatin extracts of mAC-LRR1 expressing cells synchronised in S-phase were co-immunoprecipitated with GINS antibody after treatment with p97i (5 µM) and ATRi (5 µM) for 4 hrs. Immunoprecipitated samples were analysed by western blotting with indicated antibodies.

In order to investigate this further and determine whether LRR1 needs to form a complex with CUL2 for interactions with terminated CMG on chromatin, we expressed FLAG-LRR1 or a mutant lacking the VHL box (FLAG-LRR1ΔVHL) (18), which cannot interact with CUL2 (Supp Figure 2A). FLAG-tagged recombinant LRR1 proteins (wt and ΔVHL) were immunoprecipitated from S-phase chromatin and the resulting material analysed for interactions with CMG components: MCM7, MCM2 and CDC45 (Figure 2B). Interestingly, FLAG-LRR1ΔVHL could interact with chromatin and the replisome, but its interaction with the replisomes appears somewhat destabilised (Figure 2B). We also performed a reciprocal experiment through immunoprecipitation of GINS from S-phase chromatin fraction of cells expressing FLAG-LRR1 or FLAG-LRR1ΔVHL (Figure 2C). In this case again we observed that FLAG-LRR1ΔVHL could interact with the CMG helicase. Interestingly, in both types of experiments we can observe a higher level of chromatin-bound FLAG-LRR1ΔVHL in comparison with normal FLAG-LRR1 protein, suggesting that the kinetics of interaction is affected by the lack of enzymatic activity of CUL2^LRR1^ complex.

Unfortunately, endogenous LRR1 is expressed at a very low level in cells, and we struggled to detect it reproducibly in either whole cell extracts, or in the chromatin fractions with a number of commercially available antibodies or antibodies raised by ourselves (data not shown). To detect endogenous LRR1 more efficiently, we decided therefore to use CRISPR/Cas9 to generate conditional Auxin-Inducible Degron cells using HCT116 parental cell lines, tagging endogenous LRR1 N-terminally with the mAC tag (mini AID tag fused to mCLOVER; mAC-LRR1) (Supp Figure 3A). The auxin inducible degron facilitates rapid protein degradation upon exposure to the plant hormone auxin, whilst mClover allows to enrich for tagged LRR1 protein using the GFP-trap system. First, we verified the bi-allelic gene tagging of LRR1 through genomic PCR (Supp Figure 3B) and determined that tagging of LRR1 at the N-terminus did not affect the viability of cells (Supp Figure 3C). Of note, tagging LRR1 at the C-terminus with a similar tag did not produce viable bi-allelically tagged clones, which suggested that the protein’s function is impaired (data not shown).

Taking advantage of the possibility to enrich mAC-LRR1 on GFP-trap beads, we confirmed that mAC-LRR1, expressed at endogenous levels, can interact with the replisome components and CUL2 on chromatin during S-phase (Figure 2D). We also performed the reciprocal experiment and found that immunoprecipitated GINS from S-phase chromatin could interact with CUL2 and mAC-LRR1 (Figure 2E).

### Cullin ubiquitin ligase inhibitor (CULi) traps replisomes on chromatin in S-phase and G2

Having shown that CUL2 and LRR1 interact with the replisome, we next wanted to determine whether the activity of ubiquitin ligase is required for replisome disassembly in human somatic cells. Firstly, we used the small molecule inhibitor, MLN4924 (hereafter referred to as CULi), which inhibits the neddylation pathway, primarily blocking the activation of all Cullin type ubiquitin ligases (24). It acts rapidly and does not require much previous cell manipulations to downregulate protein levels and has been used in several previous studies with model systems, including *Xenopus* egg extracts and mouse embryonic stem cells (3, 5, 6, 11, 13).

As DNA replication termination and replisome disassembly happen continuously throughout S-phase, inhibition of this process should therefore lead to higher levels of replisome components on chromatin in S-phase and prolonged retention of the replisome into G2/M stage of the cell cycle. In addition, this may potentially perturb S-phase progression if the recycling of replisome components is essential for completion of S-phase (17). General inhibition of replication fork progression (replication stress), however, can also lead to similar phenotypes, which makes it difficult to specifically distinguish between these two scenarios.

To investigate the effects of Cullin inhibition, we treated asynchronous cell cultures with a range of 1-10 µM CULi for 6 h; allowing time for cells in S-phase to progress into G2/M stage of the cell cycle so that we can observe any replisome retention. Indeed, this treatment did lead to an accumulation of MCM7 on chromatin in cells with G2/M DNA content, seen through FACS analysis in HCT116 and U2OS cell lines, with 1 µM CULi having no effect in U2OS cells (Supp Figure 4A). We then confirmed this retention of MCM7 on G2 chromatin using 5 µM CULi through FACS, immunofluorescence and western blotting of extracted nuclei (Figure 3A-C). All the above results support the role of Cullin type ubiquitin ligases in replisome disassembly in S-phase. However, cell cycle analyses also revealed an accumulation of cells with G1/early S-phase DNA content (Supp Figure 4B), reduced EdU incorporation into nascent DNA (Supp Figure 4C), fewer cells with 4N DNA content (Supp Figure 4B) and a reduction in the number of mitotic cells (Supp Figure 4D). Although CULi is very likely preventing replisome disassembly at terminating replication forks, it is important to note that it also inhibits CUL4-driven CDT1 degradation and that this stabilisation of CDT1 leads to re-loading of MCM2-7 onto chromatin, re-replication and checkpoint activation (15, 24). Indeed, we do see stabilisation of CDT1 and Cyclin E in our cells (Supp Figure 4F), as well as elevated levels of DNA synthesis (EdU incorporation) in a subset of G2 cells by immunofluorescence (Supp Figure 4G), suggesting that some G2 cells can undergo re-replication (25, 26). Considering the very high levels of chromatin-bound MCM7 visible in our FACS plots upon CULi treatment across S-phase and G2/M cells it is most likely that CULi has multiple effects: it leads to additional loading of MCM2-7 onto chromatin while impeding the unloading of active CMG helicases (Figure 3A).

**Figure 3.**
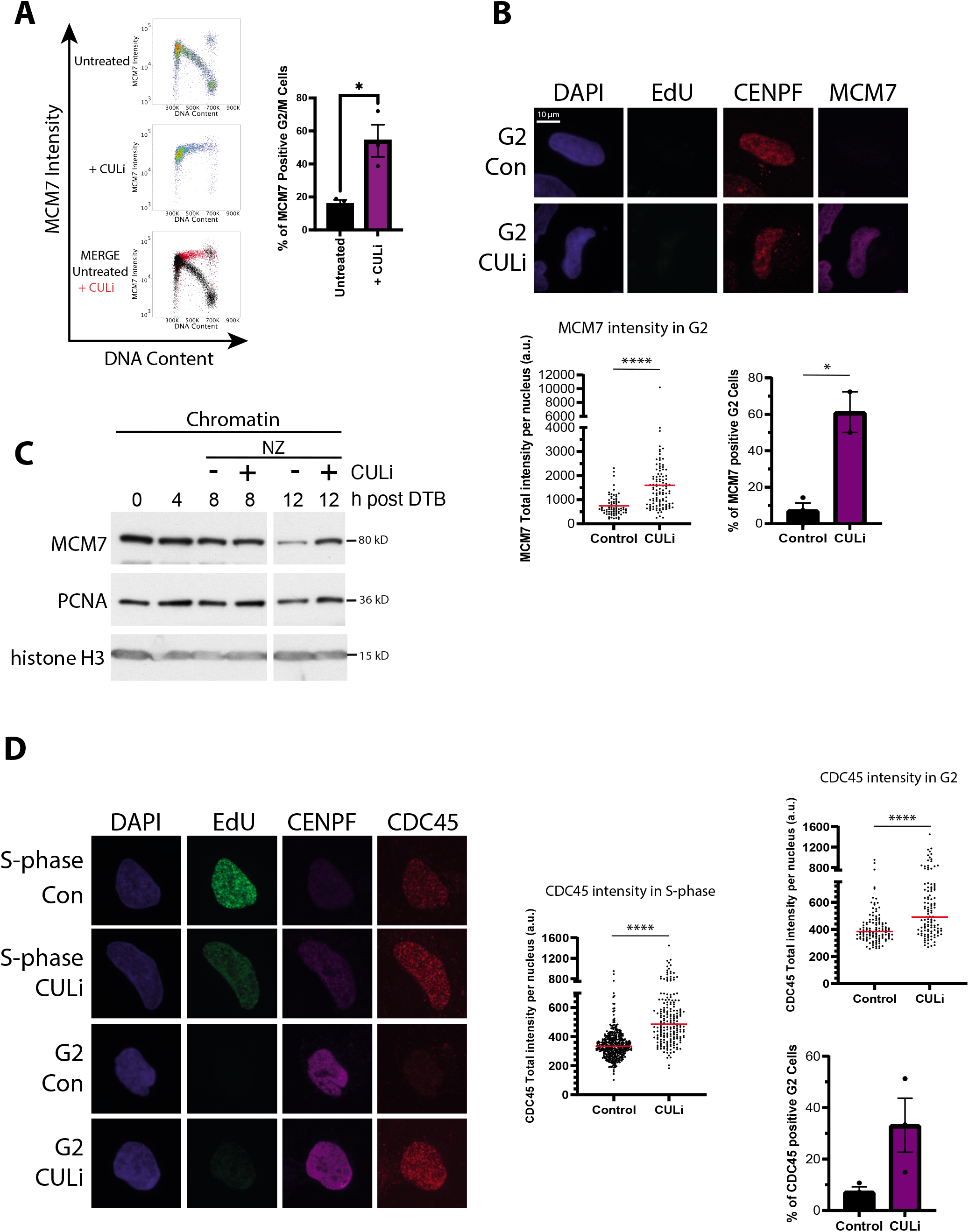
Cullin ligase inhibitor leads to replisome retention on the chromatin. **(A)** MCM7 accumulates on chromatin upon CULi treatment. Asynchronous HCT116 cells were treated with CULi for 6 h. Cells were harvested and extracted with CSK buffer to visualise MCM7 bound to chromatin. Left: example FACS plots for the total MCM7 intensity (y axis) against DNA content (x axis). Shown are untreated controls (top), + CULi (middle) and overlays of both signals (bottom). Right: Quantification of the total proportions of G2/M cells positive for MCM7 (n=3) (p=0.0187). **(B)** MCM7 accumulates on chromatin upon CULi treatment. Top: Representative immunofluorescence images of chromatin-bound EdU, CENPF and MCM7 in G2-phase U2OS cells in asynchronous population ±CULi. Bottom left: Quantification of Total MCM7 intensity in CENPF-positive/EdU-negative cells (n=3). Red lines indicate the median (p=<0.0001). Bottom right: Quantification of the total proportions of G2 cells positive for MCM7 (n=3) (p=0.0121). **(C)** MCM7 accumulates on chromatin upon CULi treatment in synchronised cells. U2OS cells were synchronised with DTB and released for indicated time points ±CULi ±nocodazole (NZ). Cells were extracted with CSK buffer and chromatin samples analysed through western blotting with indicated antibodies. **(D)** CDC45 accumulates on chromatin upon CULi treatment. Left: Representative immunofluorescence images of chromatin-bound EdU, CENPF and CDC45 in S-phase and G2-phase U2OS cells in asynchronous population ±CULi. Scatter plots: Quantification of Total CDC45 intensity in EdU-positive (S-phase) and CENPF-positive/EdU-negative (G2) cells (n=3). Red lines indicate the median (p=<0.0001 for both). Top histogram: Quantification of the total proportion of G2 cells positive for CDC45 using the high threshold (n=3) (p=0.0727); mean value with SEM.

Given this, to visualise only the active CMG complexes, and not dormant MCM2-7, we next analysed levels of chromatin-bound CDC45. Importantly, we do observe accumulation of this CMG component on chromatin in both S-phase and G2 phase cells upon CULi treatment, albeit to a lesser extent that MCM7 (Figure 3D). CDC45 is part of the active CMG helicase and is not directly re-loaded onto chromatin by stabilised CDT1. The higher levels of CDC45 on chromatin therefore likely result from combined inhibition of replisome disassembly, re-replication and inhibition of progression of S-phase by checkpoint activation. Altogether, we show that, although CULi treatment does show retainment of CMG components on chromatin in cells with S- and G2/M DNA content, it is likely an effect of wider disruptions to DNA replication during S-phase. More specific methods are therefore required for determining the function of CUL2^LRR1^ in this process.

### CUL2 depletion leads to retention of replisomes on chromatin in S-phase and G2

To target the CUL2^LRR1^ ligase more specifically, we next decided to deplete CUL2 in U2OS cells with siRNA (Figure 4A). Long term CUL2 downregulation (3 days) caused various cell cycle defects such as a reduced rate of cell proliferation (Supp Figure 5A) and a slight accumulation of cells in S-phase with FACS analysis, potentially suggesting slowed cell cycle progression (Supp Figure 5B). Furthermore, we could also detect a reduction of EdU-incorporation into nascent DNA through immunofluorescence (Supp Figure 5C), suggesting replication problems. Importantly, immunofluorescence analysis of chromatin-bound CDC45 levels in these cells revealed a significant increase in both S-phase (EdU-positive) and G2 (EdU-negative/CENPF-positive) cells (Figure 4B), suggesting that the role of CUL2 in replisome disassembly is conserved in human cells. The S-phase defects we observe could thus be a consequence of replisome disassembly defects, but to investigate this further, we decided to more specifically target the substrate receptor LRR1.

**Figure 4.**
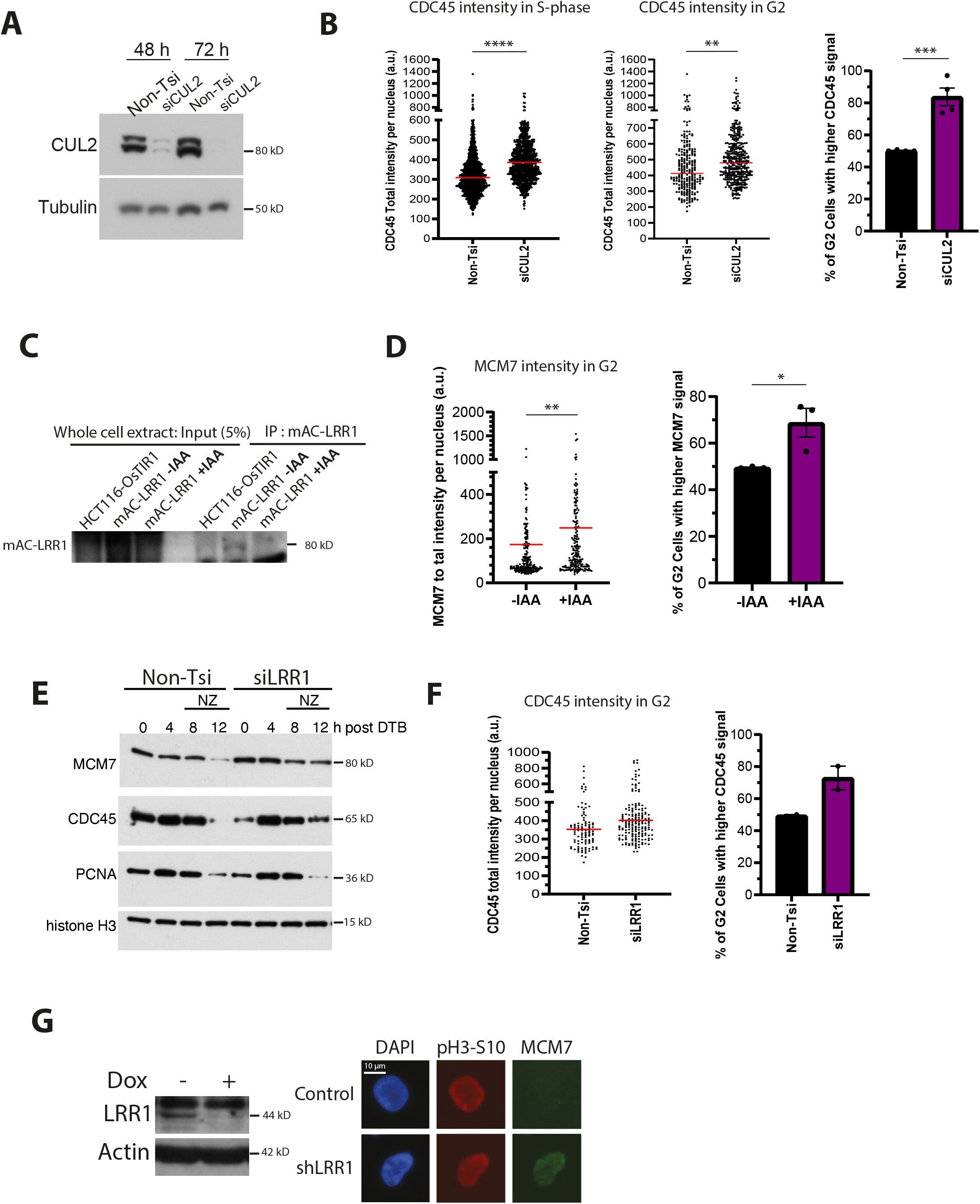
CUL2 or LRR1 depletion leads to replisome retention on the chromatin. **(A)** CUL2 downregulation by siRNA. U2OS cells were transfected with Non-T or CUL2 siRNA for 48 or 72 h. Whole cell extract samples were then analysed by western blotting with the indicated antibodies. **(B)** CDC45 accumulates on chromatin in S-phase and G2 upon depletion of CUL2. Asynchronous U2OS cells transfected with Non-T or CUL2 siRNA for 72 h were analysed by immunofluorescence. Scatter plots: Quantification of Total CDC45 intensity in EdU-positive (S-phase, left) and CENPF-positive/EdU-negative (G2, right) cells (n=4). Red lines indicate the median (S-phase p=<0.0001; G2 cells p=0.0076). Histogram: Quantification of the total proportion of G2 cells with CDC45 signal above the median of nonTsi level (n=4) (p=0.0009); mean value with SEM. **(C)** Degradation of mAC-LRR1 after 24 h of auxin treatment. Whole cell extracts of HCT116-OsTIR1 cells expressing mAC-LRR1 were lysed with UTB buffe and samples analysed by westen blotting with indicated antibodies. **(D)** MCM7 accumulates on chromatin in G2 cells upon mAC-LRR1 depletion. HCT116 LRR1-mAC cells were treated with DOX for 48 h and ±IAA for 24 h and analysed by immunofluorescence. Scatter plot: Quantification of Total MCM7 intensity in CENPF-positive/EdU-negative (G2) cells (n=3). Red lines indicate the median (p=0.0013). Histogram: Quantification of the total proportion of G2 cells with MCM7 signal above the median of -IAA cells (n=3) (p=0.0359); mean value with SEM. **(E)** CMG components accumulate on chromatin in synchronised cells upon LRR downregulation. U2OS cells were transfected with Non-T or LRR1 siRNA, synchronised with DTB and then released for indicated time points, with addition of NZ at 8 h. Chromatin was extracted with CSK buffer and samples analysed by western blotting with indicated antibodies. **(F)** CDC45 accumulates on chromatin in G2 cells upon LRR1 downregulation. Asynchronous U2OS cells transfected with Non-T or LRR1 siRNA for 72 h were analysed by immunofluorescence. Scatter plot: Quantification of Total CDC45 intensity in CENPF-positive/EdU-negative (G2) cells (n=2). Red lines indicate the median. Histogram: Quantification of the total proportion of G2 cells with CDC45 signal above the median of NonTsi control cells (n=2); mean value with SEM. **(G)** MCM7 accumulates on G2 chromatin upon LRR1 downregulation. Left: WCE of HEK293 cells inducibly expressing LRR1 shRNA for 6 days were analysed by western blotting with indicated antibodies. Right: Representative immunofluorescence images of chromatin-bound pH3-S10 and MCM7 in G2-phase cells expressing LRR1 shRNA.

### LRR1 is important for replisome disassembly during S-phase

In order to down-regulate LRR1, we made use of our HCT116 LRR1 degron cells (mAC-LRR1). Short-term knockout of LRR1 using CRISPR-Cas9 was shown previously to lead to the accumulation of CDC45 on chromatin in S-phase. To validate this in our hands, we first showed that adding indole-3-acetic acid (auxin, IAA) to the growth media indeed leads to LRR1 degradation. mAC-LRR1 was enriched on GFP-trap beads after 24 h exposure of cells to 500 µM auxin. This treatment allowed us to observe downregulation of mAC-LRR1 in the cells (Figure 4C). We then analysed the consequences of mAC-LRR1 downregulation on cell viability and cell cycle progression. Cells were grown for 10 days in the presence of doxycycline (to express OsTIR1) and 100 µM IAA, and their ability to form colonies was analysed. We observed partial reduction in the number of colonies formed in presence of IAA (Supp Figure 3D). We also analysed cell proliferation and viability, following them over a time-course of growth (Supp Figure 3E), and found partially reduced proliferation after LRR1 degradation. We then analysed the effect of mAC-LRR1 degradation on replisome unloading in S-phase and we could observe an increased level of MCM7 on chromatin in cells in G2 phase of the cell cycle (Figure 4D).

Finally, to confirm these results in other cell lines, we also downregulated LRR1 expression using si or shRNA approaches. Firstly, U2OS cells depleted of LRR1 with siRNA were synchronised in early S-phase and released. Using this approach, we found that LRR1 depletion causes a prolonged association of CMG components on chromatin through western blotting (Figure 4E), in the absence of visible changes in S-phase progression (Supp Figure 5D). In non-synchronised cells we could also detect an increase in CDC45 on chromatin in G2 cells using immunofluorescence (CENPF positive, EdU negative) (Figure 4F). Next, we generated HEK293 cells, which express an inducible pool of shRNA against LRR1, causing an 80-90% reduction in the level of LRR1 mRNA by 6 days using quantitative PCR (Supp Figure 5E). In HEK293 cells we can also sometimes detect LRR1, due to higher levels of the protein, and could therefore also detect a reduction in the steady-state levels of the protein (Figure 4G). Importantly, depletion of LRR1 with shRNA in this way also led to retainment of MCM7 on chromatin in cells with a low level of histone H3-S10 phosphorylation, representing late G2 cells (Figure 4G). Altogether, cells with depleted LRR1 have shown many fewer problems with viability, cell cycle progression and progression through S-phase, most likely reflecting the smaller pool of substrates affected by downregulation of LRR1. The lack of signs of replication stress suggests that the observed retention of CMG components on chromatin in S- and G2-stages of the cell cycle is therefore more likely an indication of problems with replisome disassembly.

### p97 inhibition (p97i) traps replisomes on chromatin but leads to many cellular problems

Next, we went on to investigate the role of p97 in replisome disassembly, with expectation that it works downstream of MCM7 ubiquitylation. To do this, we used the CB5083 inhibitor (p97i), as p97 inhibition has been shown previously to cause retention of the replisome on chromatin in model systems (5, 6, 13). In an analogous way to our studies with CULi, asynchronous cells were treated for 6 h with p97i to allow for cells to progress from S-phase into G2/M. FACS analysis of these cells revealed that the 6 h treatment with p97i led to a slightly higher level of MCM7 signal on chromatin in cells with G2/M DNA content, but the general pattern of MCM7 unloading was not much changed (Figure 5A). This suggested that the unloading of dormant origins, constituting the majority of MCM2-7 present on chromatin, is not affected by p97 inhibition. We also observed a significant increase in CDC45 on chromatin in S-phase (Figure 5B) and in G2-phase cells by immunofluorescence (Figure 5C). Altogether, these data show that while unloading of inactive Mcm2-7 double-hexamers is not affected by p97i, the unloading of CMGs is. Next, we performed the HIS pull-down with cells synchronised in early S-phase and released for 6 h in the presence of p97i. With this, we can see an increase of ubiquitylated MCM7 on chromatin, as expected from previous reports in model systems (3, 5, 6) (Figure 5D), although not as pronounced as described in embryonic model systems.

**Figure 5.**
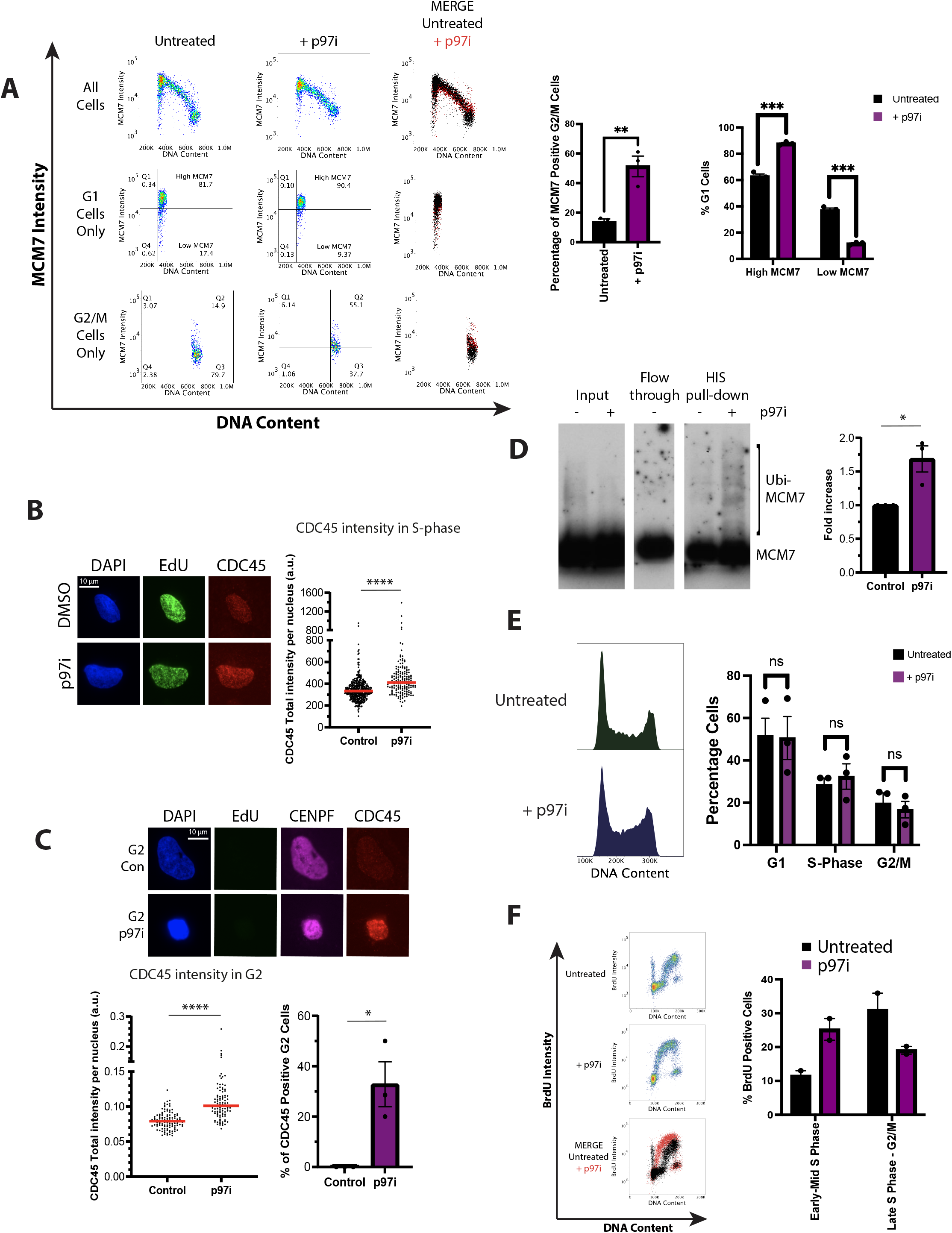
p97 inhibition leads to replisome retention on the chromatin. **(A)** p97i treatment affects loading and unloading of MCM7 from chromatin. Flow cytometric analysis of the MCM7 chromatin binding pattern following p97i treatment. Left: MCM7 levels; representative plots showing total MCM7 intensity (y axis) against DNA content (x axis). Shown is the overall binding pattern throughout the cell cycle (top), and gated examples of G1 only (middle) and G2/M only (bottom) cells used for quantification. Right: Quantification carried out over 3 repeats G2/M (p=0.0068), G1 (High MCM7: p=0.000196; Low MCM7: p=0.000171). **(B)** CDC45 accumulates on chromatin in S-phase upon p97i treatment. Left: Representative immunofluorescence images of chromatin-bound EdU and CDC45 in EdU-positive U2OS cells from asynchronous population, treated ±p97i for 6 h. Right: Quantification of Total CDC45 intensity in EdU-positive cells (n=3). Red lines indicate the median (p=<0.0001). **(C)** CDC45 accumulates on chromatin in G2 upon p97i treatment. Top: Representative immunofluorescence images of chromatin-bound CENPF and CDC45 in G2-phase U2OS cells from asynchronous population, treated ±p97i for 6 h. Scatter plot: Quantification of Total CDC45 intensity in CENPF-positive/EdU-negative (G2) cells (n=3). Red lines indicate the median (p=<0.0001). Histogram: Quantification of the total proportion of G2 cells positive for CDC45 (n=3) (p=0.0212); mean value with SEM. **(D)** Ubiquitylation of MCM7 on chromatin in S-phase following p97i. U2OS cells were transfected with HIS-Ubi plasmid, synchronised with STB and then released for 6 h ±p97i. HIS-tagged proteins were isolated using the HIS pull-down assay and samples analysed by western blotting with the indicated antibodies. Levels of MCM7 ubiquitylation were quantified using Image J. Mean with SEM (p=0.0243). **(E)** Analysis of the cell cycle profiles in asynchronous HCT116 cells following p97i treatment. Left: representative FACS plots depicting histograms of total DNA content. Right: quantification of the proportions of cells in each cell cycle stage, as described previously, n=3, (G1: p=0.945; S-phase: p=0.599; G2/M: p=0.662). **(F)** Progression of cells through S-phase upon p97i treatment. FACS analysis of HCT116 cells following the BrdU-chase experiment ±p97i for 6 h (right), with quantification (left) (n=2).

Intriguingly, the FACS profiles revealed not only a change in MCM7 chromatin binding in G2/M cells but also in G1. In the presence of p97i, in G1, we could detect only cells with fully licensed replication origins and we appeared to lose the population of cells undergoing origin licensing (Figure 5A, G1 cells). The most likely cause of this loss of MCM2-7 loading was perturbations to cell cycle progression with cells struggling to progress from mitosis back into G1, resulting in overall fewer numbers of cells loading MCM2-7 hexamers. Importantly, this cell cycle problem cannot be detected by analysing the total DNA content of cells treated with p97i for 6 h (Figure 5E), so we decided to further investigate the effects of p97i on cell cycle progression using immunofluorescence and FACS analysis. Firstly, we found a significant reduction in EdU incorporation into nascent DNA after 6 h of p97i treatment (Supp Fig 6A) (17), indicating problems during DNA synthesis in S-phase cells. Secondly, we observed a significant decrease in the total intensity of the G2 marker CENPF in all cells as well as visibly smaller G2 nuclei (Supp Figure 6B). Finally, analysis of cells positive for histone H3 S10 phosphorylation revealed clear reductions in mitotic cells upon treatment with p97i (Supp Figure 6C), as previously observed (17). These data support the idea that p97i treatment causes defects during DNA synthesis, which likely leads to checkpoint activation and fewer cells entering G2 and mitosis as a result. To test this, we turned to FACS analysis again to analyse the progression of cells through S-phase upon p97i treatment using a BrdU-chase experiment. In this, cells were treated with p97i and BrdU for 1 h, after which BrdU was washed off, while p97i remained on for another 5 h. This data clearly showed that while control cells (no p97i) can pass through S-phase and begin to re-enter G1 phase (reappearance of BrdU positive cells with G1 DNA content), cells treated with p97i struggle through S-phase (Figure 5F and Supp Figure 6D). Perturbations of p97 S-phase function in *C. elegans* embryos or human cells was previously shown to lead to S-phase checkpoint activation (25, 26). We therefore repeated the BrdU-chase experiment in the presence of p97i and the ATR inhibitor AZD6738 and found that progression through S-phase could indeed be rescued in p97i-treated cells (Supp Figure 6E). This strongly suggested that accumulating replication stress, caused by p97i-treatment, leads to checkpoint activation and cell cycle arrest, causing a reduction in the number of cells cycling through G2/M and back into G1 phase and thus observation of reduced active origin licensing. Our data suggest that p97 affects cells at every stage of the cell cycle and in result a treatment of asynchronous culture with p97 inhibitor does not show changes of overall cell cycle distribution.

Altogether, in relation to the function of p97 in replisome disassembly, we can detect higher levels of CDC45 and MCM7 on chromatin in cells with S- and G2/M DNA content and we see an increase of ubiquitylated MCM7 on chromatin in S-phase. All these are consistent with involvement of p97 in replisome disassembly in S-phase. It is important however to keep in mind that these phenotypes are likely a result of several problems during DNA replication, and not only due to a defect in replisome disassembly.

### Replisome disassembly in mitosis

Finally, given our previous studies using the *Xenopus* model system, we decided to investigate the existence of a mitotic replisome disassembly pathway in human somatic cells. As shown above, treating cells with CULi or p97i impairs cell cycle progression through S-phase and leads to reduced proportions of cells in mitosis. Despite small numbers of cells, we could see indications of MCM7 on chromatin in mitotic cells treated with p97i using immunofluorescence. These cells were identified by presenting condensed chromatin, showing a strong signal for pH3-S10 or kinetochore staining by CENPF (Figure 6A-B).

**Figure 6.**
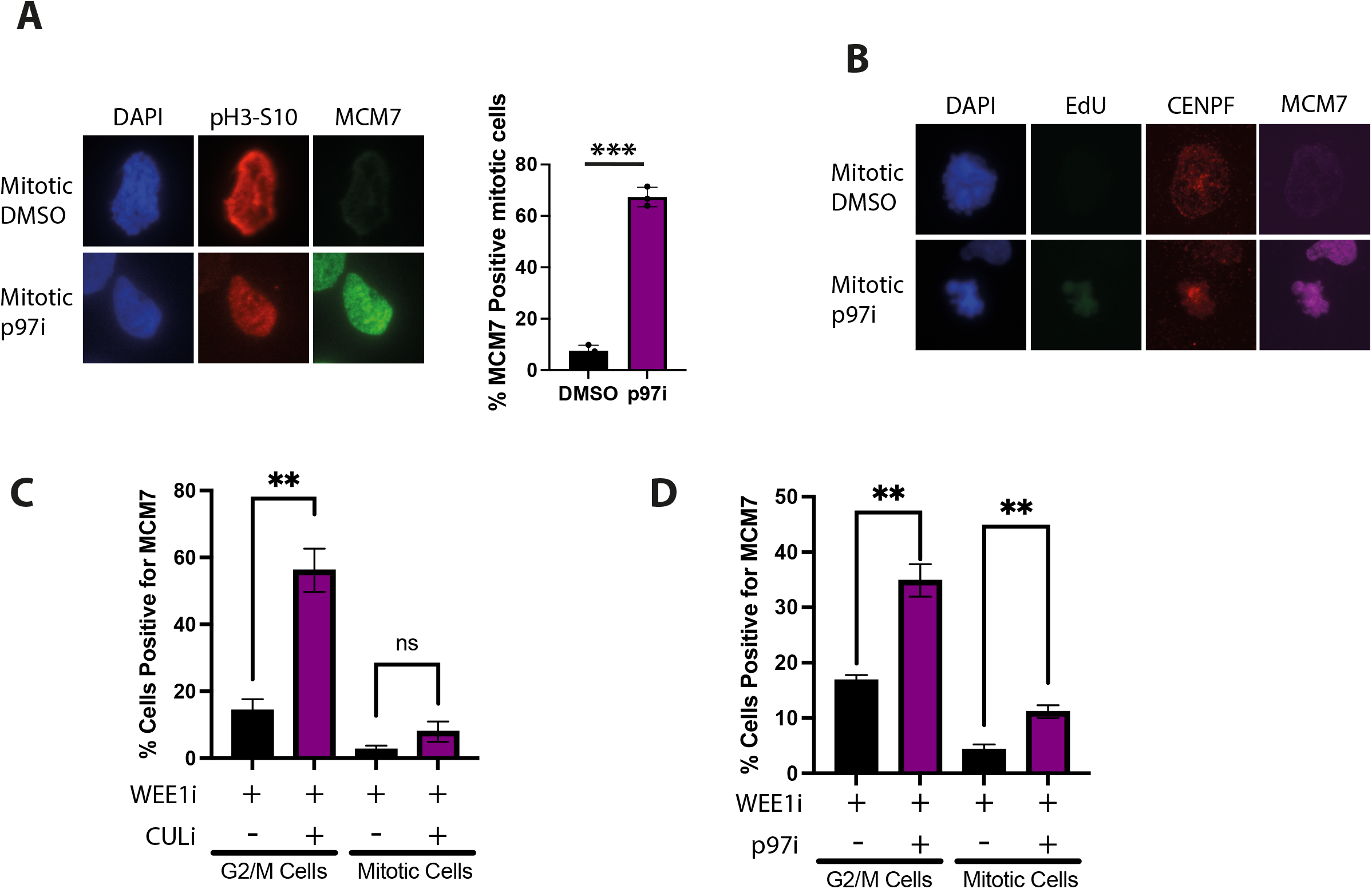
The mitotic replisome disassembly pathway is active in human somatic cells. **(A)** MCM7 is present on condensed mitotic chromosomes following p97i treatment. Left: Representative immunofluorescence images of chromatin-bound pH3-S10 and MCM7 in mitotic U2OS cells from asynchronous population, treated ±p97i for 6 h. Right: Quantification of the percentage of pH3-S10-positive cells, which are also positive for MCM7 (n=3) (p=0.000127). **(B)** MCM7 is present on condensed mitotic chromosomes following p97i treatment. Representative immunofluorescence images of chromatin-bound CENPF and MCM7 in mitotic U2OS cells from asynchronous population, treated ±p97i for 6 h. **(C)** MCM7 accumulated on chromatin following CULi treatment is unloaded from chromatin in mitosis. Analysis of the total proportion of cells retaining MCM7 in G2/M and mitosis following 6 h CULi treatment and 1 h WEE1i treatment. Cells were gated as described in Figure 2. Shown are quantification of the cells positive for MCM7 during G2/M by total DNA content (left; p=0.0045) and cells positive for MCM7 in mitosis (right, positive for pH3-S10, p=0.175). **(D)** When p97i treated cells are pushed into mitosis they still retain MCM7 on chromatin. Analysis of the total proportion of cells retaining MCM7 in G2/M and mitosis following 6 h p97i treatment and 1 h of WEE1i treatment. Shown are quantification of the cells positive for MCM7 during G2/M by total DNA content (left; p=0.0043) and cells positive for MCM7 in mitosis (right, positive for pH3-S10, p=0.0099).

To be able to visualise the mitotic pathway more quantitively, we decided to push cells treated with p97i or CULi past the G2/M checkpoint arrest and into mitosis, through inhibition of the WEE1 kinase, which deactivates CDK1 activity to prevent mitotic entry (27). First, we confirmed that a short treatment with WEE1 inhibitor (MK-1775, WEE1i) can push cells into mitosis without affecting the cell cycle or MCM7 unloading pattern (Supp Figure 7A). We next added WEE1i for the last hour of 6 h p97i/CULi treatments. Inhibition of WEE1 in CULi treated cells pushed cells into mitosis as efficiently as control cells (Supp Figure 7B). Analysis of MCM7 unloading from chromatin confirmed our previous data as there were significantly more cells with G2/M DNA content retaining MCM7 on chromatin when cells were treated with both CULi and WEE1i, but now we could also specifically analyse mitotic cells. Interestingly, although we could still detect some increase of MCM7 on chromatin in pH3-S10 positive cells, it was minor and not statistically significant (Figure 6C). Cells treated with CULi accumulate a lot of MCM2-7 on chromatin due to origin re-licensing and inhibition of replisome unloading. They do however lose these complexes from their chromatin when they enter mitosis, indicating a very efficient mitotic unloading pathway.

On the other hand, inhibition of WEE1 in p97i treated cells pushed only some of the cells into mitosis, with partial rescue of pH3-S10 positive cells numbers (approximately 50%; Supp Figure 7C). Analysis of unloading of MCM7 from chromatin revealed that there were significantly more cells with G2/M DNA content retaining MCM7 on chromatin as seen in our previous experiments (Figure 6D), but now we could detect also significantly more cells in mitosis (pH3-S10 positive cells) positive for MCM7 on chromatin (Figure 6D). This observation suggests that the mitotic replisome unloading pathway requires p97 segregase activity. Additionally, the previously described loss of G1 MCM7 loading was also rescued through WEE1i treatment, indicating that despite the persisting reduction to the proportions of mitotic populations, some cells were now able to cycle through to the G1 stage of the cell cycle (Supp Figure 7C). Altogether, these experiments suggest that the mitotic replisome disassembly pathway is indeed conserved in human cells, and p97 activity is important for unloading of the replisomes in mitosis.

## Discussion

As a process indispensable for the propagation of life, the mechanisms involved during all stages of DNA replication are well conserved throughout eukaryotic evolution. In agreement with this, we present data to suggest that the core mechanisms of replisome disassembly, first reported by our lab and others using model organisms (3-6, 9, 11, 13, 28, 29), are similarly active in human somatic cells. However, the specific characteristics of somatic cells cell cycle and S-phase regulation mean that explicit analysis of replication termination and replisome disassembly is particularly challenging. As a result, several of the tools commonly utilised for the study of DNA replication termination in model systems are less suitable for use in somatic cell models.

Replication termination and replisome disassembly events are not rare in cells. With an estimated 30-50 thousand replication origins in human cells (30), which fire during S-phase, the same number of termination events is undertaken in every cell cycle. Overall, the study of termination events is difficult when approached from a whole population level due to the highly stochastic timing and location of these events. Termination events occur whenever and wherever two progressing replication forks converge. In any individual cell a small proportion of potential replication origins become activated heterogeneously throughout S-phase, with the remaining origins (MCM2-7 double hexamers) providing a back-up in case of replication stress (31). The inactive MCM2-7 complexes, the dormant origins, are unloaded from chromatin through a yet unknown mechanism which does not require ubiquitylation of MCM7 (3). Replication forks become initiated throughout S-phase following a replication timing programme that ensures that all of the genome is fully replicated (32). As a result, the position and timing of the termination event between replication forks emanating from two adjacent origins depends on: (i) which specific MCM2-7 double hexamers were activated, (ii) timing of this activation and (iii) the speed of progression of each of the forks through the chromatin.

Moreover, replication stress experienced by the cell, either from exogenous or endogenous sources triggers a robust signalling cascade limiting replication fork progression, protecting integrity of replication forks and leading to activation of new origins around the site of damage (33). Failure to protect the replisomes at stalled forks would lead to forks collapse as the core of the replicative helicase, MCM2-7 complex, cannot be re-loaded onto chromatin during S-phase due to downregulation and inhibition of its loading factors (33). The effect of replication stress can therefore manifest itself in very similar phenotypes as inhibition of replication termination and replisome disassembly – namely accumulation of replisome components on chromatin in S-phase and into G2 stages of the cell cycle and potential slowing of progression of S-phase. At the same time, inhibition of replication fork progression (due to replication stress) limits the convergence events between forks and thus termination and replisome disassembly events. Finally, upon the replisome encountering some types of blockages on chromatin (e.g. inter-strand crosslink (ICL)), the replication machinery may also be unloaded from DNA to facilitate repair, by mechanisms independent of the canonical disassembly pathway (12, 34). To specifically focus on unloading events of terminated replisomes is therefore challenging.

In asynchronous somatic cell lines, this challenge is further exacerbated by cell cycle timing, with somatic cells possessing prolonged gap phases and only a short S-phase (35). As a result, only a small proportion of cells in a population are undergoing S-phase. To overcome this problem, cells can be synchronised to enrich for cells actively replicating DNA in the population. However, most of the synchronisation methods induce replication stress, which has to be first overcome in order for cells to resume replication. As replication stress interferes with detection of replication termination and replisome disassembly events, this solution comes with its own problems.

Due to the outlined reasons, studies to date exploring replication termination in eukaryotic cells have predominantly focused on model organisms and faster cycling embryonic stem cells, where much larger proportions of the cells actively replicate their genome (13). However, whilst particularly advantageous for the study of these processes, the use of embryonic cell lines possesses several caveats when compared to human somatic cells. Primarily, to support rapid cell division, embryonic stem cells lack several important regulatory checkpoints functioning in human cells to maintain genomic stability. Notably, these cell types often cannot induce a robust G1 or S-phase cell cycle arrest, and typically exhibit limited G2/M checkpoint activation, upon the induction of exogenous DNA damage (36, 37). Thus, it is highly likely that embryonic models are somewhat less sensitive to insults during S-phase perturbing DNA replication, that would otherwise halt cell cycle progression and ultimately cell division in somatic systems.

Starting our investigation of replisome disassembly pathways in human somatic cell lines we first confirmed that we can observe the S-phase specific ubiquitylation of MCM7 on chromatin, including with K48-linked ubiquitin chains, analogous to that reported in model systems (3, 4). The role of CUL2^LRR1^ in replisome disassembly is well described in model systems and our data established that their mechanism of action was likely conserved into human somatic cells. Immunoprecipitation of either LRR1 or components of the active replisome confirmed both Cullin2^LRR1^ complex formation and the ability to interact with the active replisome. Interestingly, we have shown that mutant LRR1 that cannot interact with CUL2 can still interact with the terminated replisome, suggesting that LRR1 does not require prior formation of the CUL2^LRR1^ complex to bind to its substrate. This is consistent with the CryoEM structure suggesting that LRR1 is the main part of CUL2^LRR1^ interacting with the replisome (7).

Following this, we compared different methods of inhibition of CUL2^LRR1^ activity to observe the effects on replisome disassembly. First, we decided to validate the use of small molecule inhibitors of all cullin ubiquitin ligases (CULi) as a tool to study this process. CULi was successfully used for this purpose both in model organisms and in mouse ES cells (3, 5, 6, 11, 13). However, although our analysis of MCM7 and CDC45 retainment on chromatin during S-phase and G2 stages of the cell cycle upon treatment with CULi is consistent with Cullin importance for replisome disassembly, it also induces DNA replication stress. CULi treatment in human cells has been reported to induce excessive genome over-replication due to MCM2-7 re-loading; through CDT1 stabilisation by the loss of Cullin4-mediated degradation (15, 24), phenotypes also found in our study. Furthermore, consistent with this, we found that the use of CULi prevented normal cell cycle progression and reduced overall DNA replication rates and mitotic cell populations.

Consequently, we asked whether the specific depletion of either Cullin2 or LRR1 would be sufficient to inhibit S-phase replisome disassembly and provide a more specific tool to block this process. We utilised RNA interference methods to deplete both Cullin2 and LRR1 in two different cell lines. In addition, auxin-inducible degradation also allowed for depletion of LRR1 by a different mechanism, targeting endogenous protein turnover instead of gene expression. In all situations, CUL2^LRR1^ depletion resulted in accumulation of MCM7 and CDC45 on chromatin in S and G2 cell cycle stages. Importantly, these downregulations led to partial inhibition of proliferation and some perturbation of S-phase progression. Analogously, short term CRISPR guided knockout of LRR1 in RPE1 cells was also shown to induce CDC45 accumulation on chromatin in late S-phase and reduce cell proliferation through defects to late S-phase progression (17), while a combination of CULi inhibitor treatment and LRR1 knockdown was required to achieve this in mouse embryonic stem cells (13). Altogether, these data indicate that CUL2^LRR1^ is indeed the ubiquitin ligase driving replisome disassembly in human somatic cell lines and the reduction of proliferation and S-phase progression problems likely represent the consequence of the loss of replisome unloading in otherwise unperturbed cell systems.

We have also investigated the involvement of p97 segregase in the process of replisome disassembly using the small molecule p97 ATPase inhibitor. Again, as with CULi, our results indicate increased retention of MCM7 and CDC45 on chromatin in S and G2 phases of the cell cycle but with an accompanying increase in levels of ubiquitylated MCM7 on chromatin. However, we do also observe induction of replication stress upon this treatment. p97 is thought to have a plethora of substrates and has been implicated in numerous mechanisms, including the DNA damage response, DNA replication, autophagy, and apoptosis (38–41). Our data indicate that a short-term p97i treatment leads to the complete loss of cell cycle progression, not initially detected through the analysis of total DNA content, as every stage of the cell cycle is affected. Importantly, cells treated with p97i cannot progress through S-phase in a timely manner. Interestingly, the delay to S-phase progression was rescued by ATRi treatment, suggesting that cells were arresting due to increased replicative stress and activation of the replication stress response. p97 with its cofactor FAF1 has been shown previously to be important for prevention of replication stress through regulation of CDT1, origin re-licensing and firing (42). This could possibly explain some of our observed phenotypes, even though the level of MCM2-7 present on chromatin upon p97 inhibition does not match the levels observed upon cullin inhibition, which leads to stimulation of origin re-licensing. Another possible path of p97 activity during S-phase could be regulation of MRE11 activity at reversed forks, as p97 was shown to prevent MRE11 retention and excessive DNA resection at double strand breaks sites (43). Finally, it has been also suggested that the loss of p97 results in vast increases to the numbers of ubiquitylated protein complexes on chromatin potentially forming barriers to replisome progression (42, 44). Altogether, p97 has a number of roles, which, when inhibited, could lead to the observed replication stress. To investigate the specific function of p97 in replisome disassembly, a more process specific tool is required. We have shown recently that the UBXN7 (UBXD7 in human) cofactor of p97 is facilitating replisome disassembly in *Xenopus laevis* egg extract (10). It will be important to test if its function is conserved in human cells.

Despite the explained above effects of small molecule inhibition to cell cycle progression, we were able to identify small numbers of mitotic cells upon p97 inhibition that retained MCM7 on chromatin. Furthermore, this population of cells was enhanced by inhibiting WEE1 kinase, which negatively regulates mitotic entry through CDK1 phosphorylation (45). This allowed us to gain an insight into the secondary mitotic, backup pathway for replisome disassembly. During mitosis, any replisomes retained on chromatin from S-phase (either terminated or stalled) are thought to be ubiquitylated by the E3 ubiquitin ligase TRAIP, resulting in their removal from chromatin by p97 (11, 28, 46). We found that treatment with CULi and WEE1i efficiently releases cells from G2 arrest and allows them to progress into mitosis. CULi leads to a large accumulation of MCM2-7 complexes on chromatin in S-phase due to re-licensing and inhibition of replisome disassembly. Nevertheless, strikingly, most of the MCM7 retained on chromatin from S-phase appeared to be disassembled in mitosis. This suggested that the backup pathway of disassembly is functional and very efficient in somatic cells. Conversely, when we combined p97i and WEE1i, only a proportion of cells managed to progress into mitosis. In these mitotic cells, replisomes were retained across both S-phase and mitosis, with a similar fold of enrichment upon p97i treatment. This is indicative of the indispensable role played by the p97 segregase in both pathways.

In conclusion, our work establishes that the two mechanisms of replisome disassembly uncovered in model organisms are indeed conserved in human somatic cell lines. We have generated tools and assays allowing us to study these processes, but also show that care needs to be taken when interpreting data as it is very challenging to specifically target these processes without other interferences.

## Materials and methods

### Cell lines

U2OS (ATCC) cell lines were grown in DMEM + 10% FBS (Sigma F7524) (Tet-free for DOX-inducible lines (PAN Biotech UK Ltd P30-3602)) + 1% pen/strep (ThermoFisher Scientific 15140122). HCT116 cell lines (ATCC) were grown in McCoys 5A media + 10% FBS (Tet-free for DOX-inducible lines) + 1% pen/strep + 2 mM L-Glutamine. HCT116 conditional degron cells were generated as detailed in (47). Bi-allelic gene tagging was confirmed using genomic PCR, see (47), using the following primers: mAID Primer ATCTTTAGGACAAGCACTCTTCTCC, LRR1 Forward: AGGAATTGCCAAGTCAAATACAG, LRR1 Reverse: AGGGAGAATATTGTGGGAGAAAG. U2OS FLAG-MCM7 and FLAG-MCM4 cells were generated using the U2OS Flp-In cell line, as previously described (48). HEK293T cells expressing tet-inducible LRR1 shRNA were generated by lentiviral infection, as detailed in (49). Briefly, for lentiviral production, low passage HEK293T cells were transfected with pTRIPZ-shRNALrr1 (Horizon Discovery) and appropriate packaging vectors (pMD2.G and psPAX2), using TransIT®-2020 Transfection Reagent (Mirus Bio 5404). Cells were cultured for 24 h before harvesting viral supernatant. U2OS cells were then transduced by centrifuging with viral supernatant containing 8 μg/ml polybrene (Sigma) for 90 min at 2500 rpm. Cells were finally selected with 1 μg/ml puromycin.

### Cell synchronisation

Double thymidine block (DTB) – exponentially growing cells were treated with 2.5 mM thymidine (Merck Life Science Ltd 6060) for 16 h to arrest them at G1/early S-phase, washed 3x with PBS, grown in fresh media for 8 h, treated a second time with 2.5 mM thymidine for 16 h, washed 3x with PBS and released into fresh media for indicated time points. Single thymidine block (STB) – exponentially growing cells were treated with 2.5 mM thymidine for 16 h to arrest them at G1/early S-phase, washed with PBS 3x and released into fresh media for indicated time points. Nocodazole (NZ) – exponentially growing cells were treated with 3.3 µM nocodazole (Fisher Scientific Ltd 15997255) for 24 h to arrest them at G2/M. Lovastatin - exponentially growing cells were treated with 20 µM Lovastatin (Acros Organics AO398529) for 24 h to arrest them at G1 phase. This is followed by 12 h treatment with 2 mM Mevalonic Acid (Sigma BCBX4250) for a synchronous release into S-phase.

### Plasmids

For generation of mAC-LRR1-degron cells, CRISPR AID plasmids were generated as detailed in (Natsume et al 2016). CRISPR-Cas9 was expressed from the pX330-U6-Chimeric_BB-CBh-hSpCas9 plasmid (50) (Addgene 42230) and targeted the N-terminus (first Methionine) of LRR1 using the complimentary guide RNA (5’ TGTAGCTTCATCTCGCCCAA 3’). Donor plasmids were also generated as detailed in (Natsume et al., 2016). Briefly, we cloned homology arms upstream and downstream of the CRISPR target region to pBluescript backbones (∼500 bp each). After inverse PCR, a cassette containing the mAC tag and hygromycin resistance marker was cloned to generate the donor plasmid. For the HIS pull-down assay, we transfected cells with pcDNA5-HIS_6_-Ubiquitin. For generation of FLAG-MCM7 and FLAG-MCM4 Flp-In cell lines, we used pcDNA5-FRT-TO-1xFLAG-MCM7, pcDNA5-FRT-TO-1xFLAG-MCM4 plasmids and the POG44 plasmid. cDNAs encoding human LRR1 were obtained from pCDNA3-FLAG-LRR1. The pCDNA3-FLAG-LRR1Δ construct was generated by using overlapping PCR mutagenesis to delete the VHL-box motif (amino acid residues 320 through 356).

### Plasmid transfection for transient expression

Cells were seeded at experiment-dependent concentrations and 24 h later they were transfected with an empirically determined optimal concentration of plasmid DNA, using Fugene HD transfection reagent (Promega UK Ltd E2312). The manufacturer’s protocol was followed and a Fugene HD transfection reagent:DNA ratio of 1:3 was used.

### FACS analysis

Following described inhibitor treatments, cells were fixed with 70% ethanol in PBS for 16 h at −20°C (typically unextracted cells or for BrdU detection), or with 4% paraformaldehyde in PBS for 15 min at RT (following CSK extractions). For antibody staining, cells were washed twice in Washing buffer (5% BSA, 0.1% Tween-20, PBS), then re-suspended in 100 µl primary antibody in Washing buffer and incubated at RT for 1 h, with rocking to prevent cells from settling. The cells were then washed twice in Washing buffer and re-suspended in 100 µl secondary antibody in Washing buffer for 1 h at RT in the dark. Stained cells were washed 1x in Washing buffer and 2x in PBS before being re-suspended in either Hoescht Staining Buffer (5 µg/ml Hoescht 33258, PBS) or Propidium Iodide Staining Buffer (50 µg/ml Propidium Iodide, 50 µg/ml RNase A, PBS). For BrdU detection, 10 µM of BrdU was added to the growth media 1 h prior to harvesting. For EdU detection, 10 µM of EdU was added to the growth media 1 h prior to harvesting. Cells were collected and fixed in ethanol as described. Fixed cells were washed once in PBS before being resuspended in 1 ml 2 M HCL supplemented with 0.1 mg/ml Pepsin for 20 min. Cells were then washed, and antibody staining carried out as described. To explore the replisome binding pattern on chromatin, cells were extracted using CSK buffer (10 mM HEPES, pH7.4, 300 mM sucrose, 100 mM NaCl, 3 mM MgCl2, 0.5% Triton X-100, 1 µg/ml of each aprotinin, leupeptin and pepstatin, 1 mM PMSF) to remove soluble fractions. The protocol used to extract cells has been described elsewhere (51). Cells were subsequently analysed with a Beckman Cytoflex instrument and data analysis performed with FlowJo (v10.7.1). Gating was performed on cell cycle profiles using the Create Gates on Peaks function.

### Immunofluorescence microscopy

Cells were seeded directly onto pre-sterilised glass cover slips, placed into individual wells of a 6-well plate at experiment-dependent concentrations. 24 h later they were treated with inhibitors or siRNA. Cells were pulse-labelled with 10 µM EdU for 20 min, before being extracted with CSK buffer (see FACS analysis method) for 10 min, to remove soluble fractions, and fixed with 4% PFA for 10 min. EdU staining was performed using the Click-It EdU kit (Invitrogen C10337) and manufacturer’s instructions, before cells were washed twice with Blocking buffer (1% BSA, 0.1% Tween-20) and incubated with the primary antibody for 1 h in the dark. After two washes with PBS and two washes with the Blocking buffer, cells were incubated with the secondary antibody for 2 h in the dark. Coverslips were then mounted onto slides using Fluoroshield + DAPI mounting medium, to co-stain for DNA. Samples were imaged using a Leica DM6000 upright widefield microscope using the 40x/1.25 Plan Apo objective.

### Immunofluorescence microscopy analysis

All images were exported as raw greyscale TIFs and analysed using CellProfiler (v4.0.6) (52, 53). The primary objects of interest (nuclei) were initially identified and masked onto the appropriate antibody staining channel. Any changes to the respective signal of individual factors within the nuclei was then determined by measuring the integrated intensity (a.u). XY scatter plots of replication factors (EdU, MCM7 or CDC45) vs CENPF were generated using Graphpad Prism (v9.2.0); examples in Supp Figure 4C. Briefly, to identify cells in G2 phase, thresholds were applied to isolate the cells that contained high CENPF intensity, whilst being negative for EdU (indicated on plots with a blue line). Any measurements to detect the intensity of CMG factors (MCM7 or CDC45) on chromatin in G2 phase were taken from cells matching the established threshold intensities only. The data was then subject to normality testing through qqplots to determine the appropriate statistical testing method (parametric or non-parametric). If parametric, an un-paired t-test was performed on the mean values of the three repeats using Prism. If non-parametric, a Mann-Whitney U-test was performed on the mean values of the three repeats using Prism. In order to quantify the proportion of G2 cells positive for the replisome factor: a positive signal threshold was determined by the highest intensity value in the control sample (excluding outliers).

However, in some experiments this approach did not reflect the differences between samples well and instead “higher CDC45/MCM7 signal” cells were quantified as ones with signal above the median value of the control cells. Statistical significance was measured by two-tailed Student’s t-test.

### Chromatin isolation

Harvested cells were washed in PBS twice before extraction in 300 µl cold CSK buffer (see FACS analysis method) for 15 min. Samples were centrifuged at 5000 rpm for 5 min, before the soluble nucleoplasm fraction was collected. Pellets were then re-suspended in another 1 ml cold CSK buffer and centrifuged again at 7000 rpm for 7 min. The final chromatin pellets were solubilised in UTB buffer (8 M Urea, 50 mM Tris-HCl pH 7.5, 150 mM β-Mercaptoethanol) and sonicated (25% amplitude – 2x 10 sec using a SONICS Vibra Cell VCX 130PB ultrasonic liquid processor). Chromatin lysates were clarified by centrifugation at max speed for 10 min and the protein concentration determined by Bradford method. Finally, they were mixed with 1x NuPAGE™ LDS sample buffer (Fisher Scientific NP0008), boiled for 5 min and analysed by western blotting as described.

### Whole cell lysate preparation

To validate LRR1 degradation in mAC-LRR1 expressing cells by western blotting, cells were lysed in UTB buffer (8 M Urea, 50 mM Tris-HCl pH 7.5, 150 mM β-Mercaptoethanol) and western blotting was performed as described. To confirm CUL2 depletion with siRNA, 7×10^5^ cells were pelleted and resuspended in 300 µl 1x RIPA buffer (50 mM Tris pH8, 150 mM NaCl, 1% Triton X-100, 0.5% sodium deoxylcholate, 0.1% SDS). Samples were vortexed every 10 min for 30 min, then sonicated (25% amplitude – 2x 10 sec using a SONICS Vibra Cell VCX 130PB ultrasonic liquid processor) and centrifuged (max speed, 10 min, 4°C). Clarified lysates were mixed with 1x NuPAGE™ LDS sample buffer, boiled for 5 min and analysed by western blotting as described.

### HIS-ubiquitin pull down

For each sample, U2OS cells were seeded into 2x 150mm dishes at 1.5×10^5^ cells/ml (30×10^5^ cells total). 24 h later they were transfected with 0.5 µg/ml pcDNA5-HIS-Ubi plasmid using Fugene HD. 32 h later, they were treated with 2.5 mM thymidine for 16 h. After 2x PBS washes, cells were treated with DMSO or p97i for 6 h, before 3×10^6^ cells were harvested for each sample (determined with a COUNTESS cell counter). Cells were extracted with CSK buffer (see FACS analysis method) + 20 mM chloroacetamide for 15 min on ice before pellets were re-suspended in 300 µl US buffer (8 M urea, 50 mM sodium phosphate, pH8) and samples boiled at 95°C for 5 min. Following sonication (25% amplitude – 3x 15 sec using a SONICS Vibra Cell VCX 130PB ultrasonic liquid processor) and centrifugation (max speed, 10 min, 4°C), 250 µl of the clarified lysate was mixed with 150 µl HIS-Tag isolation dynabeads (Invitrogen 10103D) and 750 µl US buffer for 2 h, with rotation at RT. Beads were finally washed 5x with Wash buffer (8 M urea, 50 mM Tris, pH6.8, 100 mM NaCl, 20 mM imidazole, 0.02% Tween 20, adjusted to pH6) and boiled in 2x NuPAGE™ LDS sample buffer before the original lysates (input) and the eluates (HIS pull-down) were analysed by western blotting as described.

### FLAG, GINS and GFP-TRAP immunoprecipitations

Immunoprecipitation of chromatin bound mAC-LRR1 and GINS from mAC-LRR1 expressing cells was carried out as follows. Cells were seeded at a density of 10^7^ cells/150mm dish. For efficient detection of replication termination complexes on the chromatin during S-phase by western blotting their accumulation was enhanced by supplementing the cell culture media with CB5083 (5 µM) and AZD6738 (5 µM) for 4 h before harvesting cells. Harvested cells were washed in PBS twice before extraction in cold CSK buffer (see FACS analysis method) + 20 mM chloroacetamide for 15 min. Lysates were clarified by centrifugation at 1500xg for 5 min. Chromatin pellets were solubilised in cold chromatin lysis buffer (1% NP40, 10 mM Tris pH 7.5, 100 mM KOAc, 10% glycerol, 1mM MgCl2, Universal Pierce Nuclease) followed by sonication at high amplitude for 2.5 min (30s ON 30s OFF) using a Diagenode Bioruptor Standard Sonicator. Chromatin lysates were clarified by centrifugation at max speed for 5 min followed by protein quantification by Bradford method. Where appropriate, chromatin lysates were pre-cleared by incubation with isotype control IgG coated Dynabeads at 4°C with rotation for one h. For pulling down FLAG-tagged protein, 1 mg chromatin lysates were incubated with 50 µl FLAG M2 beads (Merck A2220) for 2 h at 4°C with rotation. For pulling down GINS, 1 mg chromatin lysates were incubated with 50 µl Dynabeads Protein A (Invitrogen 10001D) coated with 10 µg GINS antibodies for 2 h at 4°C with rotation. For pulling down mAC-LRR1, 1 mg chromatin lysates were incubated with 25 µl GFP-Trap Magnetic Agarose (Chromotek) for 2 h at 4°C with rotation. This was followed by washing beads thrice with washing buffer (50mM Tris-HCl pH 7.5, 100 mM KOAc, 1 mM MgCl2, 10% Glycerol, 0.1% NP40). The immunoprecipitated protein complexes were eluted by boiling beads in 2x NuPAGE™ LDS sample Buffer at 70°C for 10 min with shaking.

### Western blotting

Proteins were resolved on a 4-12% SDS-PAGE gel and transferred onto PVDF membranes (Millipore) using Criterion™ Blotter (Bio-Rad). The membranes were then blocked using 5% milk in TBST (50 mM Tris-HCl pH 8.0, 150 mM NaCl, 0.1% Tween-20) for 1 h and incubated with appropriate primary antibodies overnight at 4°C. The membranes were washed with TBST and incubated with respective HRP-conjugated secondary antibodies for 2 h at RT. After TBST washes, membranes were developed with WesternBright ECL Spray (Advansta).

### Cell viability assays

Cells were seeded and treated with experiment-dependent conditions. To measure growth rate of cells by Trypan blue exclusion method, single cell suspensions were mixed with an equal volume of Trypan blue, and cells were counted using the automated cell counter Countess II (Thermo Fisher Scientific).

To measure growth rate by MTT assay, cells were seeded in a 96 well plate at a density of 200 cells/well and treated with experiment-dependent conditions. To measure growth rate at every 24 h, cell culture medium was replaced by medium containing Thiazolyl Blue Tetrazolium Bromide (0.5 mg/ml) (Thermo Fisher Scientific # L11939.06) and incubated for 4 h at 37°C, 5% CO_2_. Insoluble formazan salts formed by viable cell populations were dissolved by incubation in DMSO for 10 min followed by measurement of absorbance at 570 nm. Statistical significance for cell proliferation assays was measured by two-tailed Student’s t-test of unequal variance. For colony assays, asynchronous cells were diluted to a seeding density of 1×10^4^ cells/ml. Cells were seeded into individual wells of 6-well plates at the following concentrations: 250, 500, 750, 1000 total cells per well. Approximately 24 h later, cells were treated with 100 ng/ml doxycycline as appropriate, and optionally treated with 100 µM IAA 24 h later. Cells were incubated for 7-10 days, or until sufficient colonies were observed, replacing the media every 3-4 days. At this point, the cell media was removed, and cells stained with methylene blue (2% methylene blue in 50% ethanol) for 5 min at RT and the plates dried at RT overnight. Colonies were counted and first normalised to the seeding density to obtain the percentage colony forming units. Per condition, each respective % colony forming units calculated were then normalised to untreated controls (Tet-OsTIR1 untreated) to allow for comparison between cell lines and seeding densities. Data were analysed using one-way ANOVA, with any statistical differences detected subsequently explored using post-hoc testing (TukeysHSD) using RStudio. Colony assay data was graphed using RStudio, with the packages ggplot and ggpubr.

### siRNA transfection

Cells were seeded at 5×10^4^ cells/ml and 24 h later transfected with 50 nM Non-targeting (Non-T, Horizon Discovery D-001810-10-05), CUL2 (Horizon Discovery D-007277-01-0005 and D-007277-03-0005) or LRR1 (Horizon Discovery L-016820-01-0005) siRNA using Dharmafect 1 (Horizon Discovery T-2001-02), following manufacturer’s protocol.

### qPCR

qPCR was performed as described in (49) using LRR1 primers forward: GGGACCCGCTATGAGCTAAG and reverse: CCTTTAACCGAACAGTGGCTTT.

### Inhibitors and reagents

5 µM CULi (MLN4924, Stratech Scientific S7109-SEL. Dissolved in DMSO to generate 5 mM stock); 5 µM p97i (CB5083, Generon Ltd B6032. Dissolved in DMSO to generate 5 mM stock); 5-10 µM ATRi (AZD6738, Generon Ltd B6007. Dissolved in DMSO to generate 10 mM stock); 2 µg/ml DOX (doxycycline hyclate, VWR International Ltd CAYM14422. Dissolved in DMSO to generate 2 mg/ml stock); 100 µM IAA (3-Indole Acetic Acid, Sigma I2886. Dissolved in DMSO to generate a 500 mM stock); 5 µM Wee1i (MK-1775, Cayman Chemicals, 21266. Dissolved in DMSO to generate a 10 mM stock); 1 µg/ml puromycin (ThermoFisher Scientific A1113803. Dissolved in HEPES buffer; 10 mg/ml stock); 100 µg/ml hygromycin B GOLD (ThermoFisher Scientific 10687010. Dissolved in H_2_O; 50 mg/ml stock).

### Antibodies

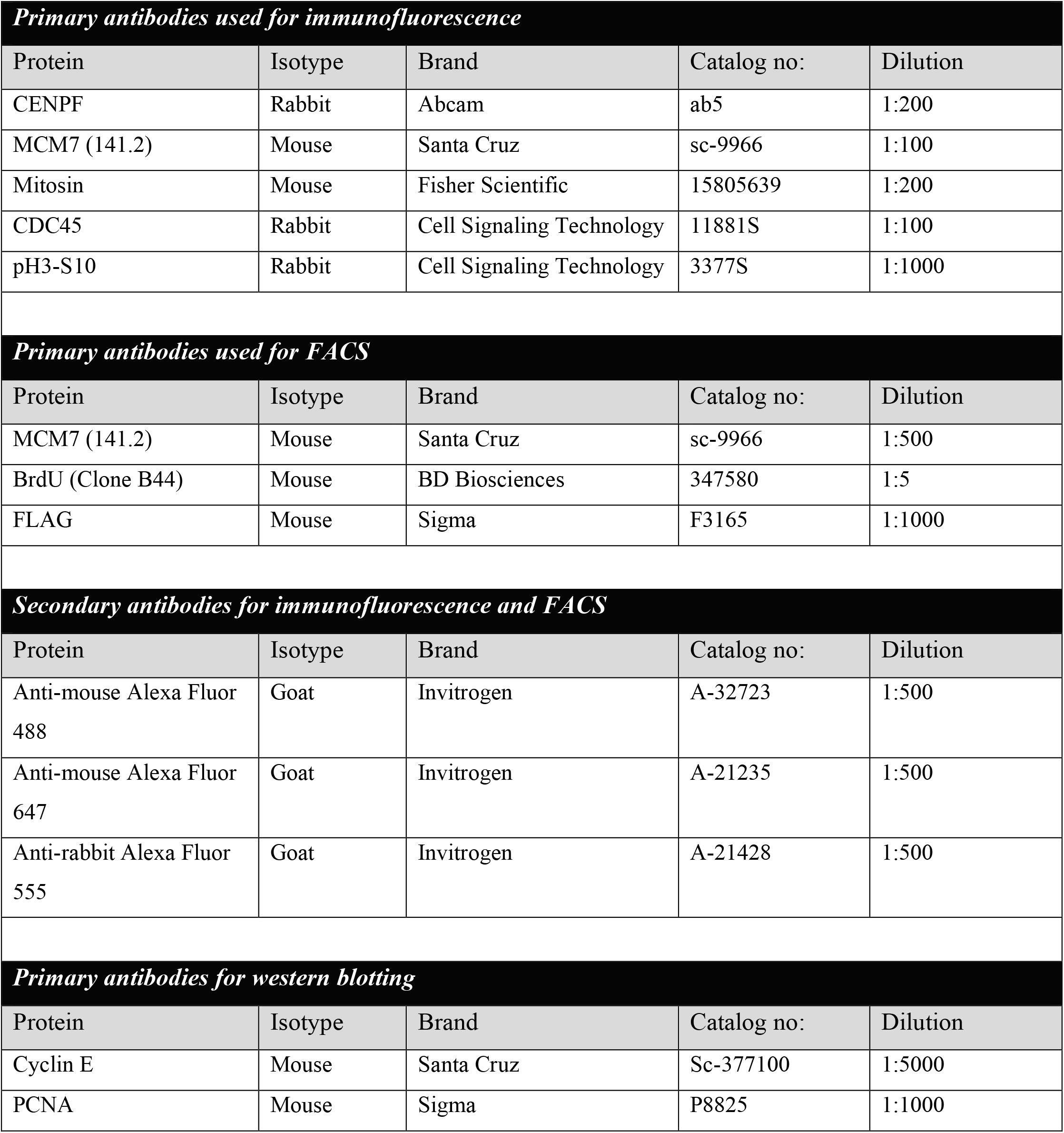
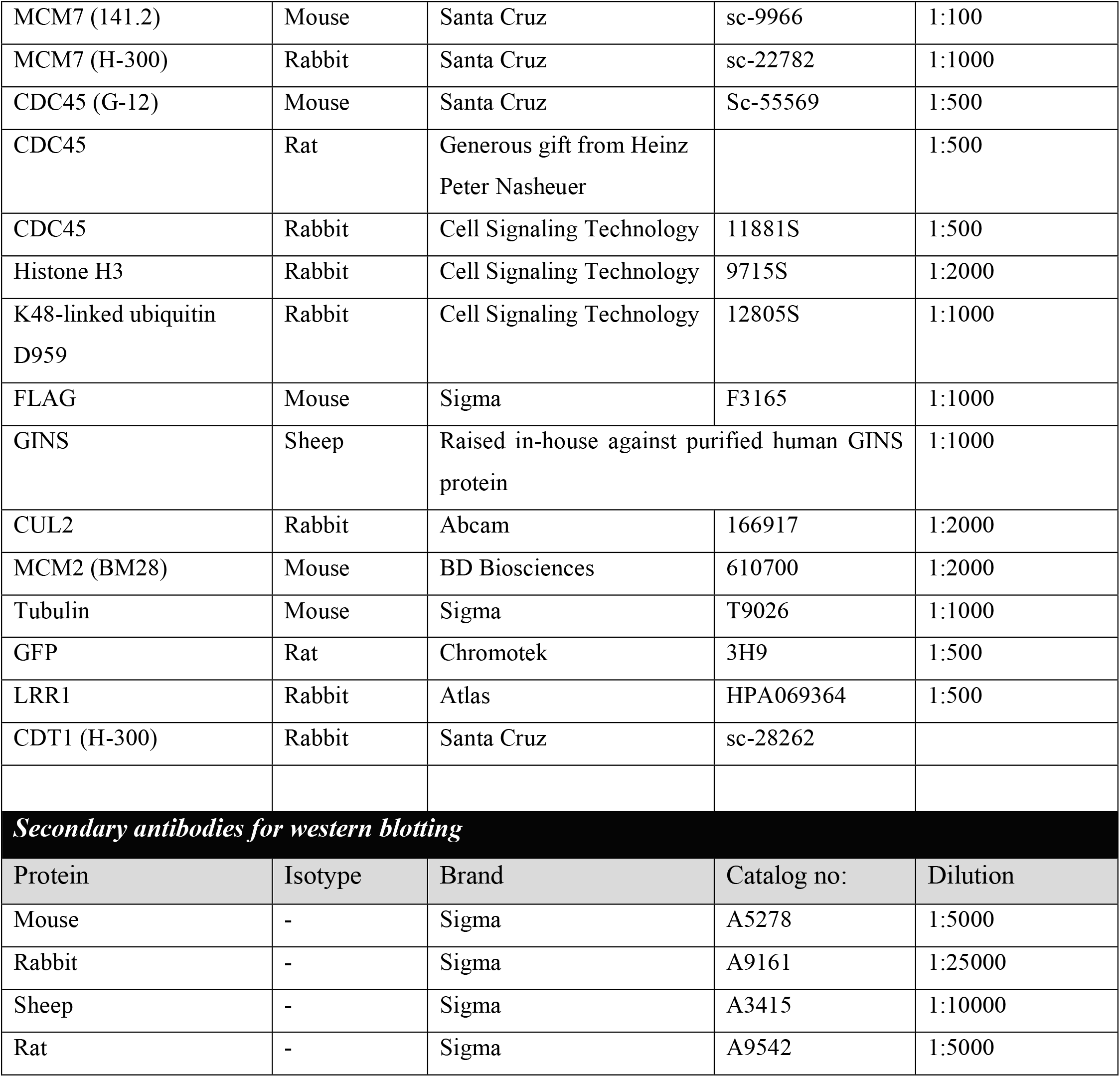

## Supporting information

combined supplementary figures

## Author Contributions

R.M.J., J.H.R., S.S., S.N. and M.H. generated and analysed the data; T.N. and M.K. assisted with generation of degron cell lines; R.J., S.S. and A.G. wrote the manuscript; A.G. designed and overlooked the project.

## Acknowledgments

This work was supported by the Lister Award and Wellcome Trust Investigator Award (215510/Z/19/Z) for A.G., BBSRC funded MIBTP studentship and JSPS Summer programme for S.S. and University of Birmingham. We would like to thank Dr Neville Gilhooly and Dr Marco Saponaro for critical discussions of the manuscript.

## Notes

### Competing Interest Statement

The authors have declared no competing interest.

