## Supplementary material for "Characterising replisome disassembly in human cells": combined supplementary figures

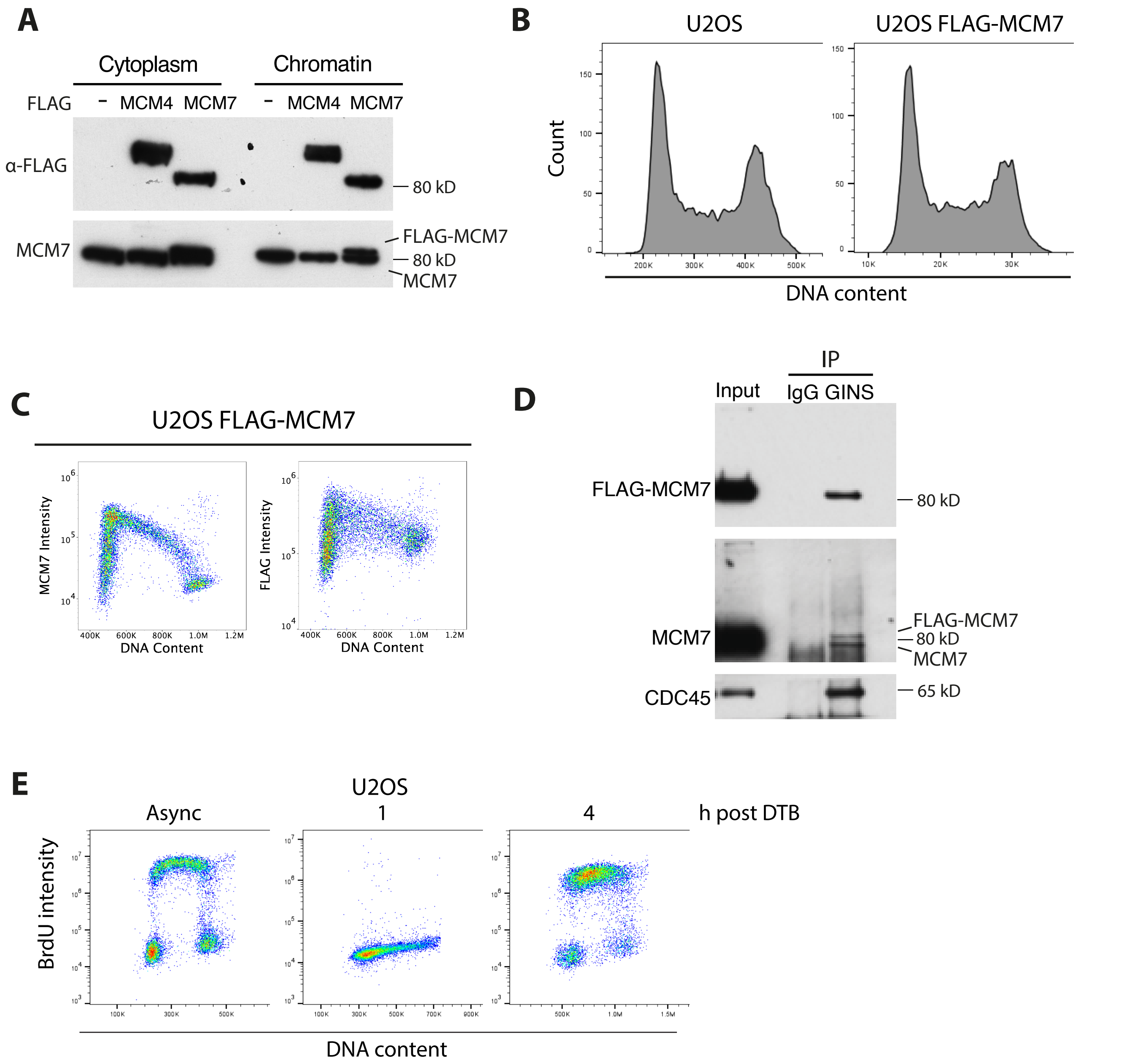


**Supplementary Figure 1. Tagging MCM7 with FLAG-tag does not affect its function. (A)** Cytoplasmic or chromatin fractions were extracted from U2OS cells expressing FLAG-MCM4 or FLAG-MCM7 and samples analysed by western blotting with indicated antibodies. **(B)** Cell cycle profiles for U2OS and FLAG-MCM7-expressing U2OS cells. **(C)** MCM7 chromatin binding pattern of U2OS cells expressing FLAG-MCM7. Representative data shown are FACS plots from the same sample, with MCM7 and FLAG intensity measured in different channels. **(D)** U2OS cells expressing FLAG-MCM7 were synchronised in S-phase with DTB and GINS was immunoprecipitated from chromatin lysates.
**(E)** U2OS cells synchronised with DTB, released for indicated time points and pulsed with BrdU for 1 hour prior to harvesting. BrdU intensity and DNA content measured in each sample by FACS.

Jones et al. Supplementary Figure 2

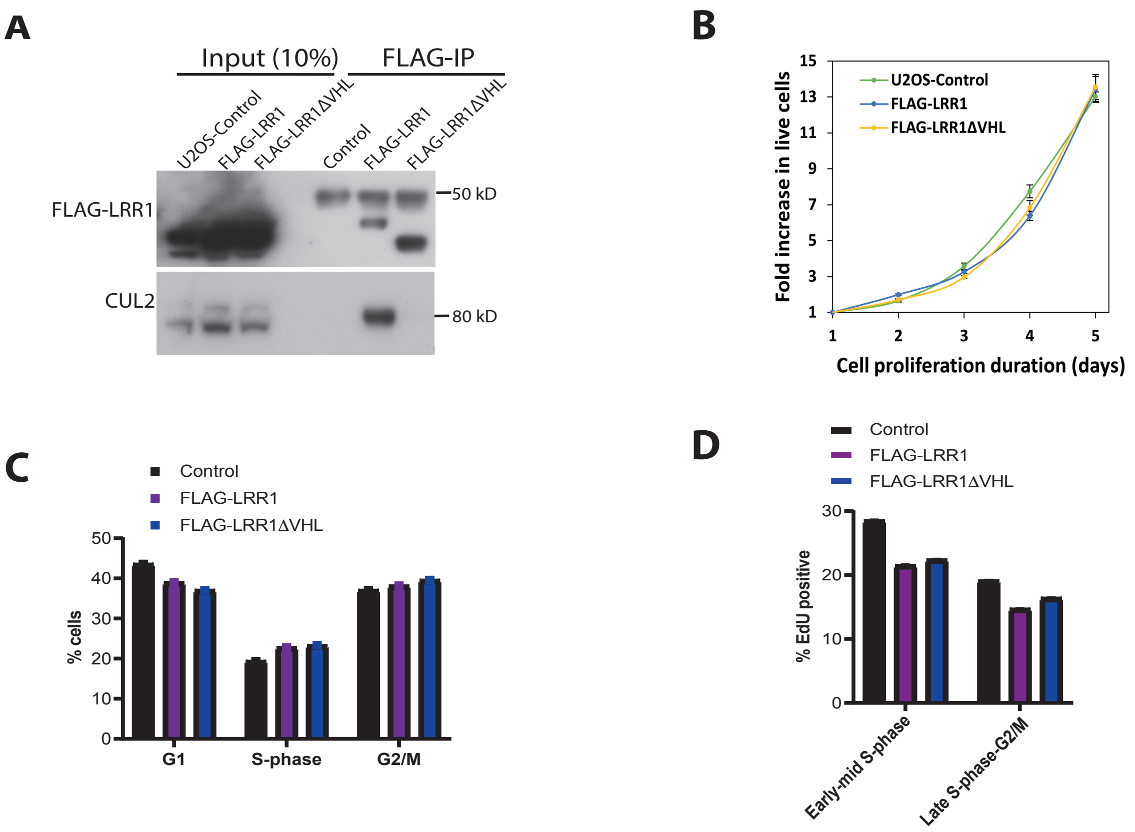


**Supplementary Figure 2. Expression of FLAG-LRR1 or FLAG-LRR1ΔVHL does not affect cell proliferation.** **(A)** FLAG-LRR1 or FLAG-LRR1ΔVHL were immunoprecipitated from whole cell extracts of U2OS cells with FLAG M2 beads and analysed with antibodies against the indicated proteins. **(B)** Fold increase of live U2OS cells expressing FLAG-LRR1 or FLAG-LRR1ΔVHL (n=3). **(C)** Analysis of the cell cycle profiles of control U2OS cells, or U2OS cells transiently transfected with FLAG-LRR1 or FLAG-LRR1ΔVHL plasmids for 48 h. **(D)** Quantification of EdU FACS analysis in control U2OS cells, or U2OS cells transiently transfected with FLAG-LRR1 or FLAG-LRR1ΔVHL plasmids for 48 h.

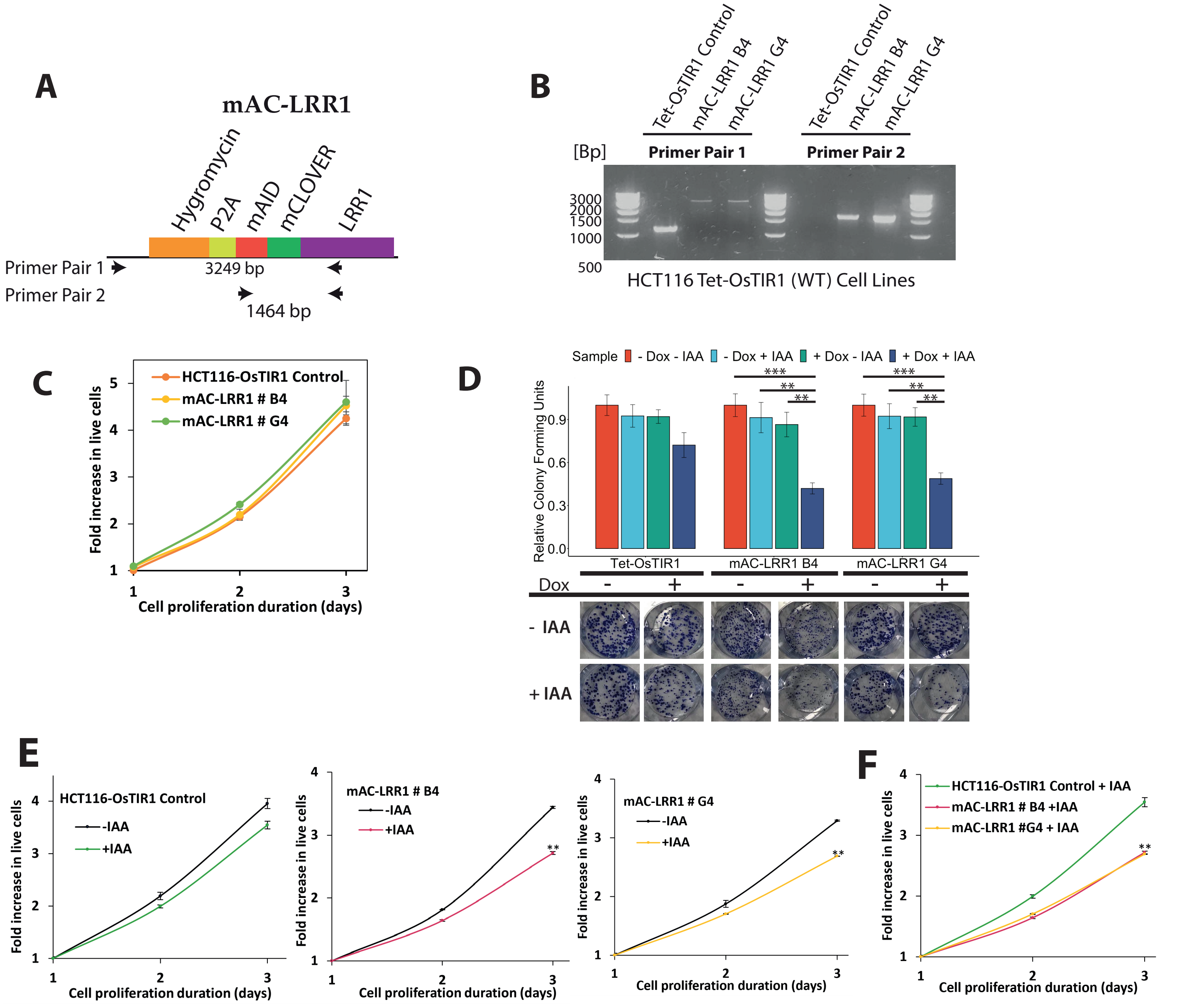


**Supplementary Figure 3. Characterisation of HCT116 cells with IAA-induced LRR1 degrons.**
**(A)** Tagging layout for mAC-LRR1 degron cells. Endogenous LRR1 was tagged N-terminally with a degron tag, p2a self-cleavage site, and hygromycin resistance marker. Arrows indicate the position of primers deigned to screen for bi-allelic gene tagging. Expected sizes resulting from the amplification of these regions is also shown. **(B)** Example DNA agarose gel following genomic PCR to screen for cells in HCT116 Tet-OsTIR1 (WT) background. Shown are the confirmed bi-allelic cell lines utilised for this study. **(C)** Fold increase in live HCT116 cells expressing mAC-LRR1 was monitored for 3 days. n = 3; error bars indicate standard deviation (SD). **(D)** Colony assays showing cell viability for the selected HCT116 mAC-LRR1 cells. Colonies were counted (n = 3) and normalised to untreated controls (top). One-way ANOVA: Tet-OsTir1 p=0.0749; mAC-LRR1 B4 p=0.000274; mAC-LRR1 G4 , p=0.000154). Pairwise hypothesis testing within the degron cells (Tukeys HSD), with significant differences shown on the plot. **(E)** HCT116 cells expressing mAC-LRR1 were treated with Tetracycline (1µg/ml) and IAA (100 µM) and fold increase in live cells was monitored for 3 days. n=3; error bars indicate standard deviation (SD) *p=<0.05; **p=<0.01. **(F)** As (E) but auxin (IAA) treated control cell line and mAC-LRR1 degron cell lines are compared.

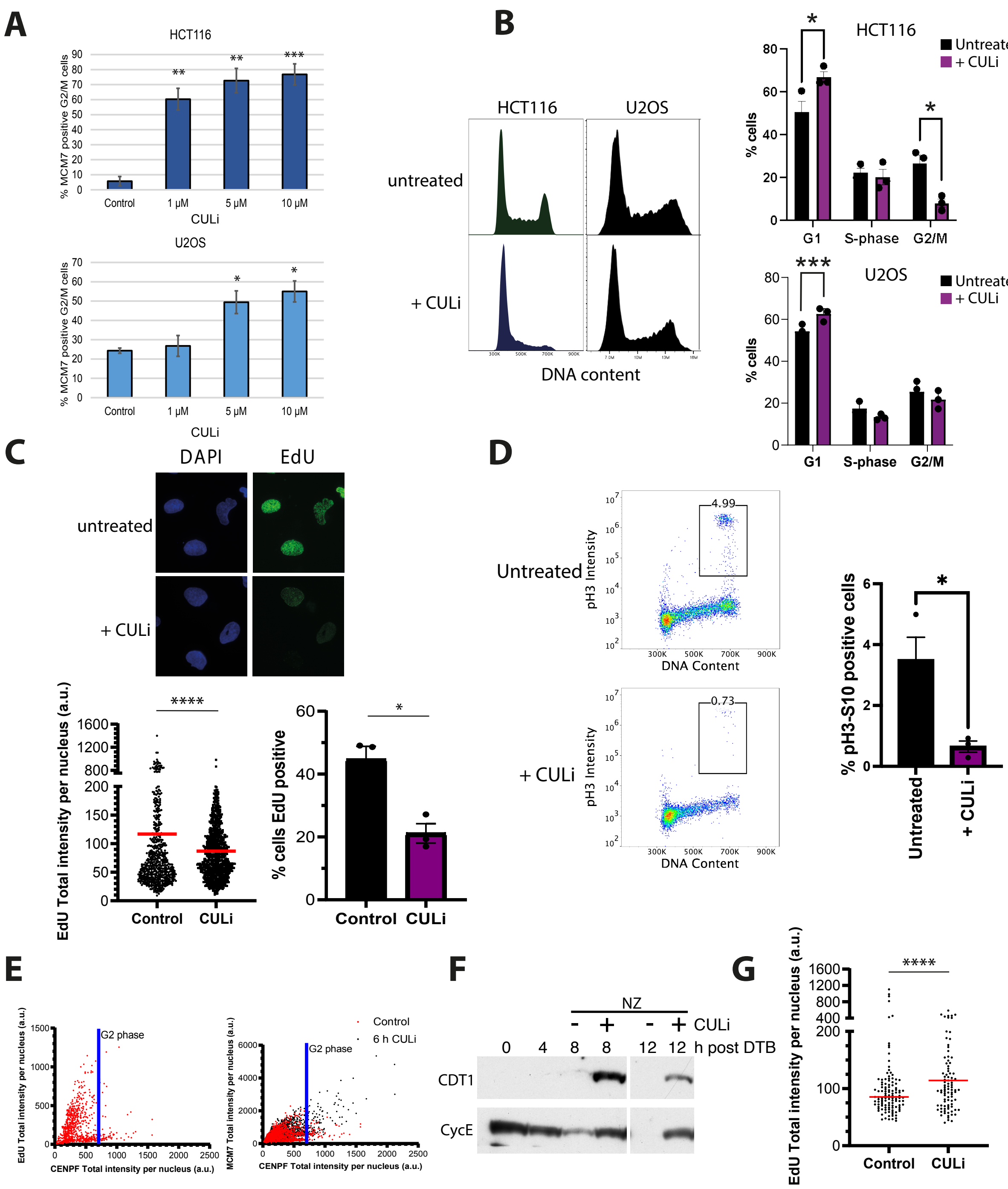


**Supplementary Figure 4. The effects of cullin activity inhibition. (A)** FACS analysis quantification of % of G2/M cells positive for MCM7 from asynchronous HCT116 and U2OS cells, treated with differing concentrations of CULi for 6 hours (n=3). HCT116: 1 µM (p=0.00199), 5 µM (p=0.0019), 10 µM (p=0.00085). U2OS: 5 µM (p=0.0261), 10 µM (p=0.0372). **(B)** CULi affects cells progression through S-phase. Cell cycle profiles of asynchronous HCT116 and U2OS cell lines following optional CULi treatment for 6 h. Left: Representative histograms depicting total DNA content (x axis). Right: Quantification of the proportion of cells in G1 (2N DNA content), S-phase (2N < DNA Content > 4N), or G2/M (4N DNA content) over 3 experimental repeats. (HCT116: G1: p=0.0434; G2/M: p=0.0134. U2OS: G1: p=0.0004. **(C)** CULi affects cells ability to synthesise DNA. Top: Representative immunofluore-scence images of EdU in asynchronous U2OS cells, treated ±CULi for 6 h. Scatter plot: Quantification of EdU total intensity in all cells (n=3). Red lines indicate the median (p=<0.0001). Histogram: Mean with SEM (p=0.0103): **(D)** Analysis of the propotion of mitotic (pH3-S10-positive) cells following 6 hours CULi treatment. Left: example FACS plots depicting pH3-S10 intensity (y axis) against total DNA content (x axis). Right: Associated quantification over 3 repeats (p=0.0212). **(E)** See Microscopy analysis in Materials and Methods. Left: Example XY scatter plot of EdU vs CENPF in asynchronous population of cells. Right: Example XY scatter plot of MCM7 vs CENPF in asynchronous population of cells. The threshold is shown for marking G2 cells, as described in Materials and Methods. **(F)** U2OS cells were synchronised with DTB and released for indicated time points ±CULi ±nocodazole (NZ). Chromatin s amples analysed through western blotting with indicated antibodies. **(G)** Quantification of EdU total intensity in G2 cells from Figure 2F (n=3). Red lines indicate the median (p=<0.0001).

Jones et al Supplementary Figure 5

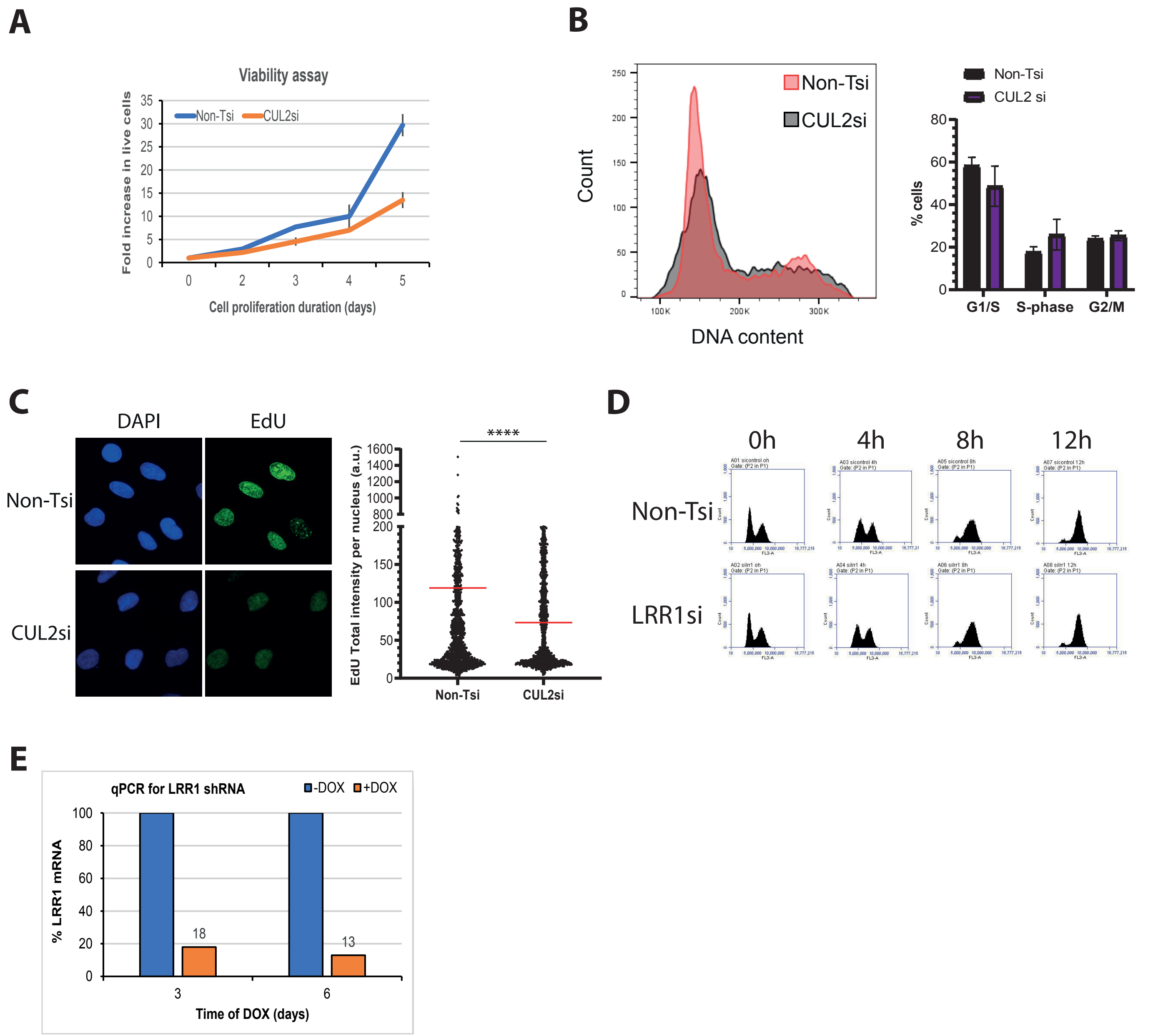


**Supplementary Figure 5. Downregulation of CUL2 leads to loss of viability and DNA replication problems.** **(A)** Growth assay to compare proliferation of U2OS cells with Non-T or CUL2 siRNA. Cells were transfected on day 1 (n=1 (duplicate)). **(B)** Cell cycle profile of U2OS cells depleted of CUL2 with siRNA for 3 days (n=2). **(C)** Left: Representative images of EdU-positive U2OS cells, from asynchronous population following transfection with Non-T or CUL2 siRNA for 3 days. Right: Quantification of total EdU intensities (n=3). Red lines indicate the median (p=<0.0001). **(D)** U2OS cells depleted of LRR1 with siRNA were synchronised with DTB and released for indicated time points and cell cycle profiles analysed by FACS. **(E)** qPCR analysis for LRR1 mRNA in HEK293 cells stably expressing DOX-inducible LRR1 shRNA for 3 or 6 days.

Jones et al. Supplementary Figure 6

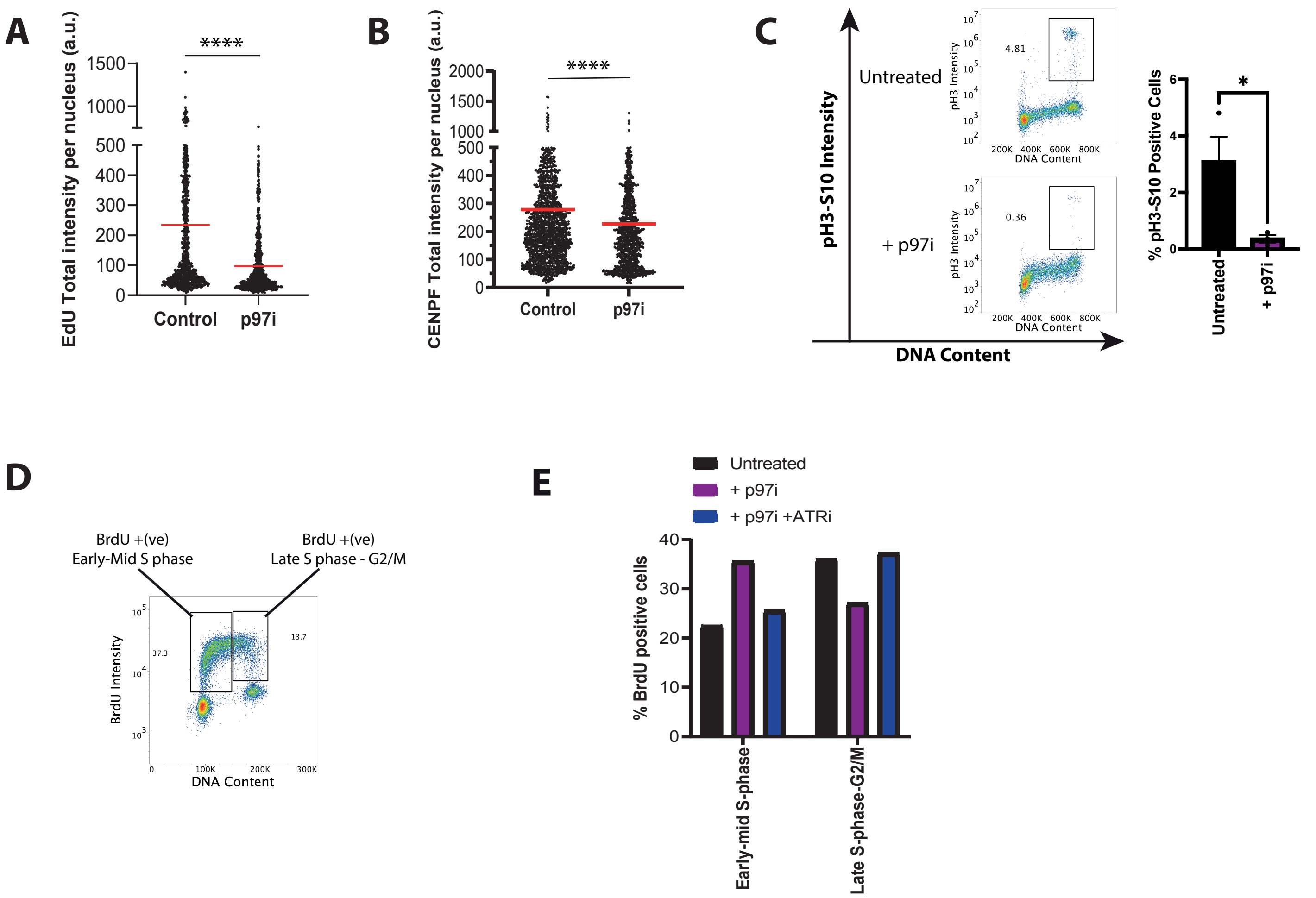


**Supplementary Figure 6. The effects of p97 activity inhibition. (A)** Quantification of total EdU intensity in all cells treated ±p97i for 6 hours (n=3). Red lines indicate the mean (p=<0.0001). **(B)** Quantification of total CENPF intensity in all cells treated ±p97i for 6 hours (n=3). Red lines indicate the mean (p=<0.0001). **(C)** Analysis of the total proportion of mitotic cells (pH3-S10 positive) upon p97i treatment. Left: Representative plots showing pH3-S10 intensity (y axis) against total DNA content (x axis). Black boxes indicate mitotic cell gates. Right: quantification of the total proportions of cells positive for pH3-S10 (p = 0.0369). **(D)** Gating example for Figure 5F. Shown is a representative plot depicting the gating strategy used. Gates were established using control samples (no BrdU chase) and applied to all test samples. **(E)** Slow S-phase progression in p97i treated cells is due to checkpoint activation. Quantification of FACS analysis of U2OS cells following the BrdU-chase experiment ±p97i ±ATRi for 6.5 h.

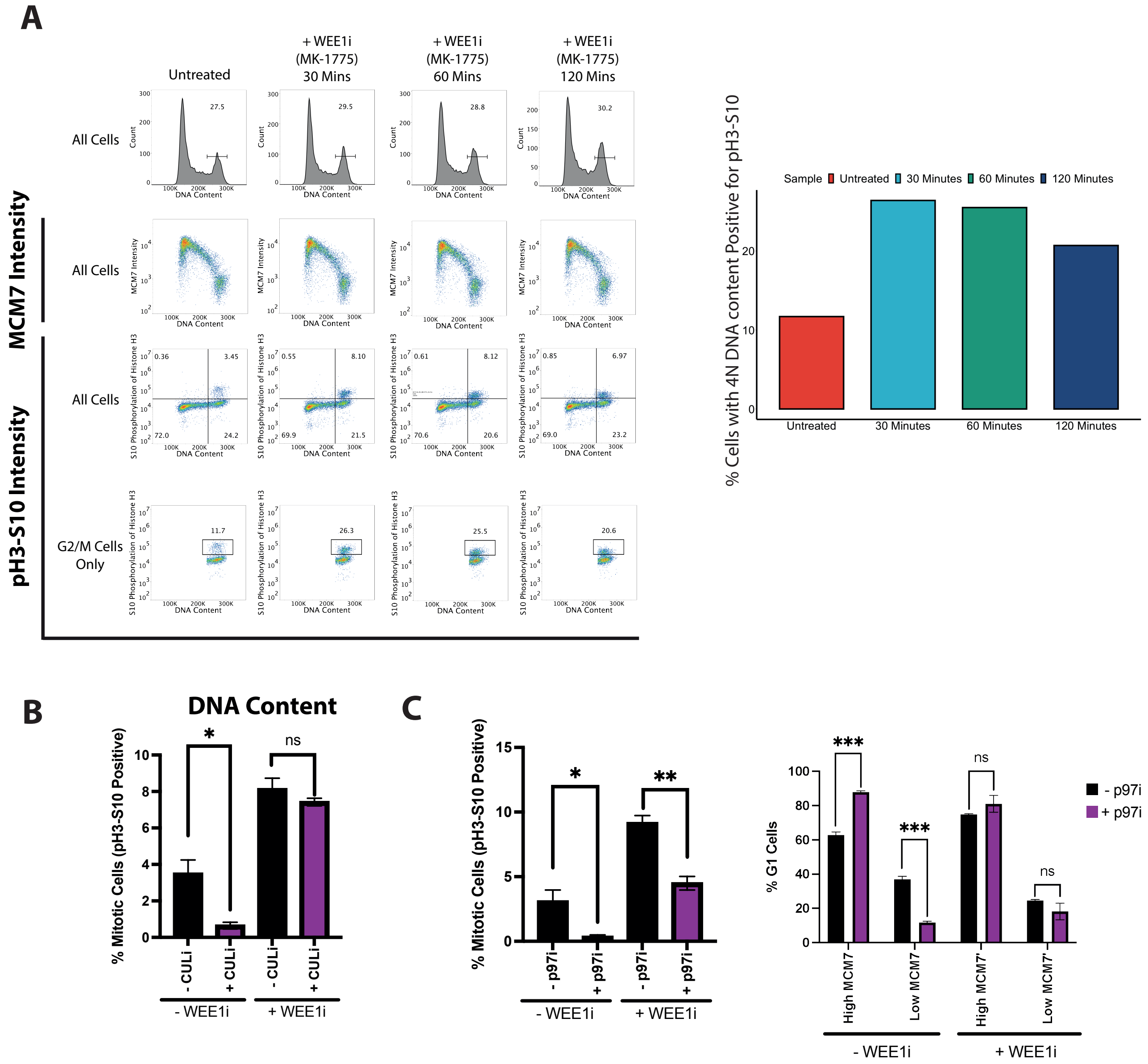


**Supplementary Figure 7. Inhibition of WEE1 kinase allows CULi or p97i treated cells to enter mitosis.** **(A)** Validation of the ability of WEE1i to push cells into mitosis. Asynchronous HCT116 cells were treated for the described time with WEE1i. Left: Representative FACS plots showing the overall cell cycle, MCM7 binding pattern, pH3-S10 staining pattern, and mitotic cells. Right: Quantification of the proportion of mitotic cells (4N DNA cells positive for pH3-S10). **(B)** Effects of CULi in combination with WEE1i on cell cycle progression. Cells were treated for 6 hours with CULi, supplemented for the final hour with WEE1i. Left: quantification of the proportion of mitotic cells. **(C)** Effects of p97i in combination with WEE1i on cell cycle progression. Cells treated as in (B). Left: quantification of the proportion of mitotic cells (t.test: $p = 0.0103$). Right: quantification of the approximate amounts of cells undergoing origin licensing.
